# Virus-Driven Proximity Proteomics Identifies p400 as a Restriction Factor Poised on Incoming Viral Genomes

**DOI:** 10.64898/2026.08.17.745356

**Authors:** Tanner M. Tessier, Katelyn M. MacNeil, Orlando B. Scudero, Jack W. Dowling, Mackenzie J. Dodge, Cason R. King, Joseph S. Mymryk, Matthew D. Weitzman

**Affiliations:** Division of Protective Immunity, Children’s Hospital of Philadelphia, Philadelphia, USA; Department of Pathology and Laboratory Medicine, Perelman School of Medicine, University of Pennsylvania, Philadelphia, USA; Department of Microbiology & Immunology, University of Western Ontario, London, ON, Canada; Cell & Molecular Biology Graduate Group, University of Pennsylvania Perelman School of Medicine, Philadelphia, PA, USA; Department of Medical Microbiology & Immunology, University of Wisconsin-Madison, Madison, USA; Department of Otolaryngology, University of Western Ontario, London, ON, Canada; Department of Oncology, University of Western Ontario, London, ON, Canada; Penn Epigenetics Institute, University of Pennsylvania Perelman School of Medicine, Philadelphia, PA, USA

## Abstract

Immediate-early viral proteins must rapidly counteract host defenses directed at incoming viral genomes, yet the protein interaction networks that mediate this remain unresolved. This gap persists largely because immediate-early proteins are expressed at low abundance during this stage of infection, making them challenging to investigate with standard approaches. Here, we present a virus-driven proximity proteomics framework to investigate immediate-early phase virus-host interactions in an authentic infection context. An isogenic P2A control virus untethers a miniTurbo biotin ligase from the viral protein under investigation, effectively modeling bait abundance and background changes from a matched infection context. Continuous biotin labeling from the earliest hours of infection further amplifies detection of transient, low-abundance interactions characteristic of this phase. Using the adenovirus E1A hub protein as a benchmark, we recovered the majority of known E1A interactors and identified over 150 high-confidence interactions. Applying this strategy to the immediate-early phase, we resolved the E1A interactome and identified the p400 chromatin remodeling complex as its dominant target. We show p400 associates with incoming viral genomes, represses gene expression, and regulates persistent infection. These data identify p400 as a poised host restriction factor and establish a temporally resolved proximity proteomics strategy transferable to other immediate-early viral proteins.

## Introduction

Virus- and host-driven protein-protein interactions (PPIs) play central roles in shaping the viral replication cycle and the cellular defenses that restrict infection^1,2^. By integrating into host PPI networks, viral proteins hijack and influence nearly all aspects of the normal cellular program^3–6^. Detailed analysis of virus-host PPI networks therefore remains fundamental to understanding the molecular basis of viral replication and host biology and has direct implications for identifying druggable host factors or pathways to control infection^7,8^.

The viral replication cycle is classically divided into immediate-early, early, and late phases. The immediate-early phase is arguably the most consequential, encompassing both the initiation of viral gene expression and the subversion of host defenses directed at incoming viral genomes. The balance between these opposing processes determines whether an abortive or productive infection is established. Viruses have therefore evolved elegant strategies, carried out by their immediate-early proteins, to navigate this window. Critically, strategies to identify the virus-host PPIs that govern this phase, with accurate spatial and temporal resolution, remain largely unaddressed. Established approaches such as AP-MS^9,10^, thermal proximity coaggregation^11^, and crosslinking mass spectrometry^12^ rely on abundant viral protein expression and an already-remodeled host proteome, neither of which is present during the earliest hours of infection. As a result, nearly all virus-host PPI studies to date have been carried out at time points beyond the immediate-early phase, leaving this stage of infection mechanistically underexplored.

Proximity labeling has emerged as a powerful and complementary approach to AP-MS, with the additional advantage of having the sensitivity to capture weak and transient PPIs^13^. Proximity labeling has been used to study virus-host interactions through transient transfection^14–17^ and others have incorporated biotin ligases into the virus to interrogate interactions in an infection-relevant context^18–20^. Despite the value of these studies, they share important limitations relevant to mapping PPIs with temporal precision in an infection context. Most importantly, neither employ a stoichiometry-matched, infection-context control. Instead, background is typically defined by mock infection, an unfused ligase, or empty vector, none of which account for the dynamic changes in bait protein abundance and the host proteome itself are over the course of infection. Therefore, these approaches are susceptible to conflating changes in PPI stoichiometry with changes in bait abundance or infection-driven proteome remodeling.

To address these limitations, we designed a proximity proteomics strategy built around a paired isogenic control virus containing a P2A ribosomal skipping sequence to untether miniTurbo^21^ from the bait of interest. This design allows bait abundance and infection-driven background proteome changes to be modeled directly from a virus expressing the same set of proteins under authentic infection kinetics, rather than inferred from a mismatched control. We further reasoned that the catalytic nature of biotin ligation could be leveraged to overcome the low bait abundance that would typically preclude interactome mapping during the immediate-early phase. By initiating labeling immediately upon infection and allowing biotin to accumulate over the immediate-early phase, transient early interactions are cumulatively biotinylated and their “signal” amplified. To benchmark this approach, we selected the adenovirus hub protein E1A, a small, intrinsically disordered protein that uses short linear motifs to engage a disproportionately large PPI network relative to its size^22,23^. E1A is the first protein expressed during infection and is an essential regulator of the immediate early phase^24–26^, making it an ideal test for our proteomics approach. Using our paired viruses, we recovered the majority of known E1A interactors and identified numerous novel putative interactors, many derived from large multi-subunit transcription and chromatin-remodeling complexes. Applying this strategy specifically to the immediate-early phase, we identified the p400 chromatin remodeling complex as the dominant E1A target and found that p400 associates with incoming viral genomes to attenuate early viral gene expression. We additionally show p400 regulates persistent infection, which remains poorly understood despite the life-threating clinical outcomes for immunocompromised and hematopoietic cell transplantation populations^27,28^. Together, this work establishes a stoichiometry-controlled, temporally resolved strategy for resolving virus-host PPI networks during transient and technically challenging stages of infection, and identifies p400 as a constitutively poised host restriction factor.

## Results

### A Paired P2A Control Virus Enables Stringent Detection of E1A-Proximal Interactors

We engineered wild-type human adenovirus C5 (HAdV-C5) to encode a FLAG-tagged miniTurbo (FmT) biotin ligase fused directly to the N-terminus of E1A (FmT-E1A) (**Fig. 1A and B**). MiniTurbo was selected for its small size, high biotinylation activity, and strict dependence on exogenous biotin, which together allow tight control over a defined labeling window^21^. Because the N-terminus is shared across all E1A splice isoforms, this design ensures each isoform is tagged with FmT. E1A abundance increases continuously over the course of infection, and this increase coincides with widespread infection-driven remodeling of the host proteome^24,29–33^, making it difficult to stringently identify high-confidence PPIs against a stable proteome baseline. To control for these factors simultaneously, we generated an isogenic control virus, FmT-2A, identical to FmT-E1A except for a P2A ribosomal skipping sequence inserted within the linker separating FmT and E1A. Untethered FmT expressed from this virus is present at stoichiometric levels that track FmT-E1A expression as its expression is controlled by the same viral genomic elements. However, it biotinylates proteins non-specifically given that it is not covalently fused to E1A. Additionally, infection kinetics and the accompanying remodeling of the host cell proteome should progress similarly between the two viruses. This novel control allows high-confidence E1A interactors to be resolved against an infection-matched background, rather than an uninfected or mismatched control that does not adequately reflect viral reprogramming of the infected cell.

**Figure 1.**
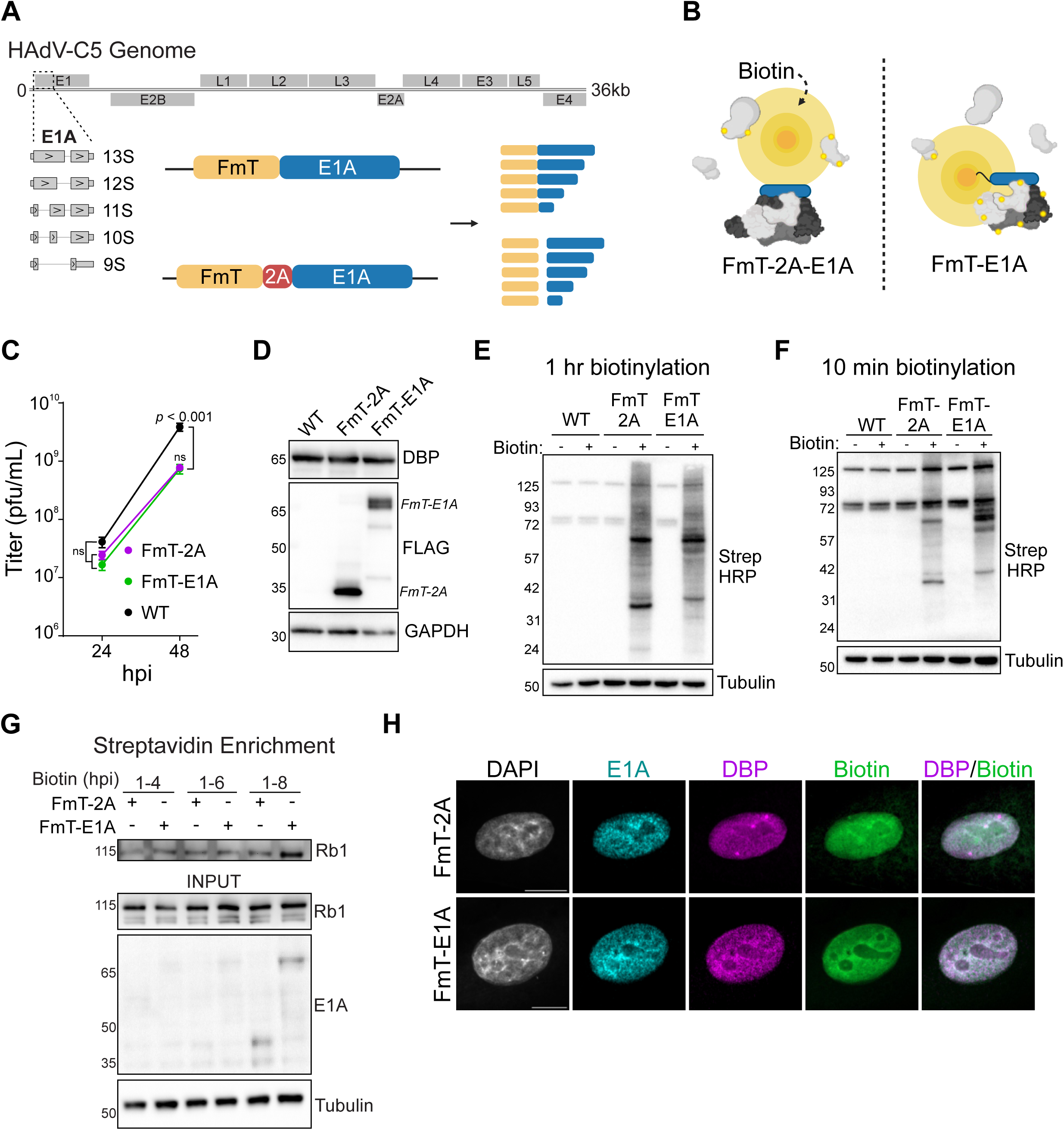
Virus-encoded proximity-proteomics. **A)** Flag-miniTurbo (FmT) virus schematic. FmT-E1A and FmT-2A-E1A are expressed from endogenous regulatory elements within a wild-type HAdV-C5 genome. FmT-2A-E1A contains a P2A ribosome skipping sequence. A schematic of the HAdV-C5 genome and E1A transcripts are depicted. N-terminal FmT will tag all E1A isoforms. **B)** Untethered FmT-2A will reflect FmT-E1A expression and will non-specifically biotinylate proteins within the infected cell. Comparison to FmT-E1A will allow for determination of high confidence E1A protein interactors. **C)** Infectious progeny production in A549 cells infected with WT HAdV-C5 or FmT viruses. Titer was determined by plaque assay on HEK 293 cells. Two-way ANOVA with Tukey’s correction, n = 3. **D)** Anti-FLAG detection of FmT-2A and FmT-E1A in infected A549 cells at 24 hpi. DBP is used as a marker of infection. **E and F)** Biotinylated protein in whole cell lysates from infected A549 cells. Biotinylation was added for 1 hour or 10 minutes at 24 hpi. **G)** Streptavidin enrichment of biotinylated FmT-E1A interactors during the immediate early phase of infection. Biotin was added from 1-4, -6, or -8 hpi and biotinylated proteins were enriched. Rb1 is used as a representative E1A interactor. **H)** Confocal immunofluorescence microscopy in A549 cells infected with FmT viruses at at 24 hpi. Biotin was added for 30 minutes to label E1A interactors. DBP is used to mark viral replication compartments. Scale = 10 µm.

We first confirmed that our modified viruses preserve wild-type infection kinetics. Both FmT-E1A and the FmT-2A control virus exhibited productive splicing and expressed both viral early and late proteins at levels similar to wild-type HAdV-C5 over a time course of infection (**Fig. S1A**). Late protein expression and DBP were modestly decreased in both FmT virus infections, but by 32hpi they were expressed at levels similar to wild-type unmodified HAdV-C5 (**Fig. S1A**). While lower than wild-type infection, infectious progeny production was identical between FmT viruses (**Fig. 1C**). Western blotting for FmT confirmed efficient P2A cleavage and closely matched levels of tethered and untethered FmT (**Fig. 1D**). Together, these data demonstrate that our P2A design strategy functions as a stoichiometry-matched background control, while both viruses show identical infection kinetics that closely approximate wild-type infection.

We then characterized the labeling behavior of the FmT viruses. MiniTurbo exhibited minimal background activity in the presence of endogenous biotin, but rapidly biotinylated proximal proteins upon addition of exogenous biotin (**Fig. 1E**). Robust labeling was also detectable after as little as 10 minutes (**Fig. 1F**). Background biotinylation remained remarkably low even at 48 hours post-infection (hpi), when accumulation of endogenous background would be most apparent (**Fig. S1B**). We next determined if we could enrich for known E1A interactors within the first 8 hours of infection, when E1A is barely detectable by western blot. Following infection with our FmT viruses, we initiated biotinylation at 1 hpi and allowed this to proceed until 4, 6 and 8 hpi, where we then enriched for biotinylated proteins via streptavidin pulldown (**Fig. 1G**). By 8 hpi, we could confirm enrichment of Rb, a well-established E1A interactor^34^, in FmT-E1A infected cells. Together, these data demonstrate our approach retains sensitivity and specificity even when bait abundance is minimal.

Confocal immunofluorescence microscopy in A549 cells showed that E1A and biotinylated proteins were exclusively nuclear at both early and late time points during FmT-E1A infection, indicating that FmT tagging does not perturb E1A nuclear localization (**Fig. 1H, S1C, and S1D**). During FmT-2A infection, biotinylation remained predominantly nuclear with only minor cytoplasmic signal, consistent with FmT dissociating from E1A in this control virus (**Fig. S1D**). Collectively, these data establish that FmT-2A provides a stoichiometry-matched, infection-context background for identifying E1A-proximal interactors. Moreover, rapid and low background biotinylation enables the temporal and spatial precision needed to resolve interactions at defined stages of the viral replication cycle.

### A Combined DDA/DIA Proteomics Workflow Identifies High-Confidence E1A Interactors Across Infection Conditions

To demonstrate that our approach can identify virus-host PPIs in an infection context, we mapped the E1A interactome at typical time points when E1A is abundant and host remodeling is established, as well as under conditions of activated innate antiviral immunity (**Fig. 3A**). A549 cells were infected with FmT-E1A or FmT-2A virus and biotinylated from early (15–17 hpi) or late (31–33 hpi) time points, or from 15–17 hpi following 16 hours of IFNβ pretreatment. To maximize protein identification and reproducibility, we combined DDA and DIA mass spectrometry, which together identified up to 2,500 proteins across all samples with low missing-value rates (2–8% per sample) (**Fig. S2A**). Principal component analysis showed discrete clustering by condition/timepoint (PC1, 40%) and viral infection type (PC2, 18%) (**Fig. S2B**), and the majority of proteins (>85%) were quantified in all replicates per condition (**Fig. S2C**), indicating a robust dataset for comparing FmT-E1A directly against the FmT-2A control.

Comparing high-confidence PPIs across conditions identified 150–190 interactors per condition, which was not entirely surprising given that E1A is a well-recognized hub protein with many reported interactions^22,23^ (**Fig. 2A**, Supplemental Data 1). Many interactors were shared across conditions, though each also possessed a unique subset (**Fig. 2B**). Since FmT-2A tracks FmT-E1A abundance, fold-changes for shared interactors can be compared directly across time points despite differences in absolute E1A abundance. Many shared interactors showed similar fold-changes relative to FmT-2A across conditions, suggesting these PPIs scale with E1A abundance rather than reflecting condition-specific regulation (**Fig. 2C**). Consistent with E1A biology, at 16 hpi we see preferential interactions with Rb family members, RBL1 and RBL2, which are essential E1A targets for dysregulating cell cycle early during infection. Other PPIs, such as MAX and NRIP1, were preferentially enriched under IFNβ stimulation, pointing to putative interactions specifically favored during an antiviral response. Together, these data show that our paired-virus control strategy resolves both abundance-scaled and condition-specific components of the E1A interactome within a single framework during viral infection, while also identified novel interactors.

**Figure 2.**
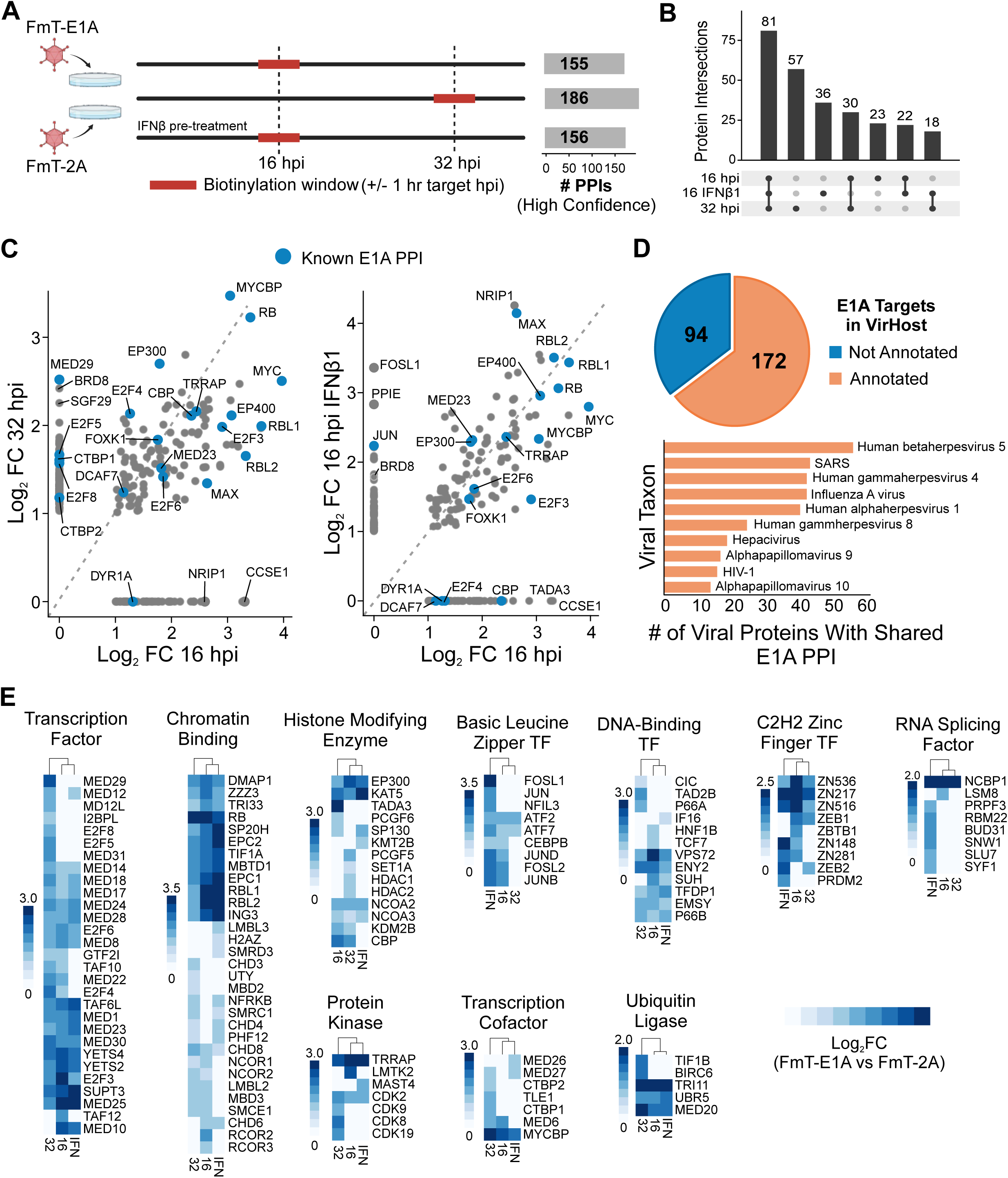
Characterization of the E1A proximity interactome during infection. **A)** Schematic of proteomics experimental design to determine high confidence E1A proximal interactions during infection in A549 cells. The number of high confidence interactors (adj.*p* val ≤ 0.05 and log2 fold-change ≥ 1) from each condition are indicated. n = 3. **B)** Upset plot of overlapping high-confidence interactors. **C)** Comparison of high confidence interactors based on their FmT-2A normalized log2 fold-change. Examples of known E1A interactors are labeled in blue. **D)** Comparison of all high confidence E1A interactors with virus-host PPIs annotated in the VirHost database. Other viral taxa with proteins that also targets E1A interactors identified in our experiment are displayed below. **E)** Top 10 PANTHER protein families based on high-confidence PPIs from each condition.

### The E1A Interactome is Dominated by Transcriptional Regulators and Unannotated Virus-Host Interactions

Among the putative novel E1A interactors, we also recovered numerous established ones, including Rb^34^, E2F^35^, p300/CBP^36,37^, p400^38^, MYC^39,40^, CTBP1/2^41–43^, and Mediator complex subunits^44,45^ (**Fig. 2C and S3A**). CBP/p300 and p400 are both targeted through the E1A N-terminus^38,46^, CTBP1 and 2 bind the E1A C-terminus ^43^, and Mediator binds centrally within conserved region 3^44,45^, collectively demonstrating our approach captures interactors spanning the full length of E1A and is unaffected by the FmT fusion. Under IFNβ stimulation, we observed specific enrichment of the AP-1 transcription factors FOSL1 and JUN, which are associated with stress-induced interferon signaling^47,48^. Notably, these interactors have previously been linked to E1A only by *in vitro* binding assays^49^.

To determine whether these interactors extend beyond the established E1A interactome, we compared all 266 high-confidence PPIs identified across conditions to the VirHost database of curated virus-host PPIs^50^. Notably, approximately one-third of E1A interactors in our dataset are not annotated in VirHost (**Fig. 2D**), and those that are annotated are shared across multiple viral taxa. Based on PANTHER classification^51^, transcription factors and chromatin-binding proteins represented the largest category of high-confidence PPIs, alongside smaller groups of histone-modifying enzymes and leucine-zipper and zinc-finger transcription factors (**Fig. 2E**, Supplemental Data 2). In fact, approximately one-third of all interactors are classified as either a transcription factor or transcription cofactor^52^ (**Fig. S3B**). Additionally, several protein classes, particularly RNA splicing factors and leucine-zipper transcription factors, were enriched predominantly within the IFNβ condition. These classifications are consistent with E1A’s established role as a major transcriptional activator^53,54^, while substantially expanding the repertoire of host factors implicated in E1A-mediated transcriptional rewiring.

### High-Confidence E1A PPIs Map Onto Host Protein Complexes

To characterize the structural organization of the E1A proximity interactome, we analyzed high-confidence interactors for membership in known protein complexes using the CORUM and Complex Portal databases^55,56^ (**Fig. 3**, Supplemental Data 3). Many E1A interactors mapped onto large multi-subunit complexes, several of which have not previously been linked to E1A, including the mutually exclusive MBD2- and MBD3-containing NuRD nucleosome remodeling complexes^57^. These data also provide more complete evidence for interactions with complexes where only a few E1A interactors have been identified in the past, such as the SAGA complex^39^.

**Figure 3.**
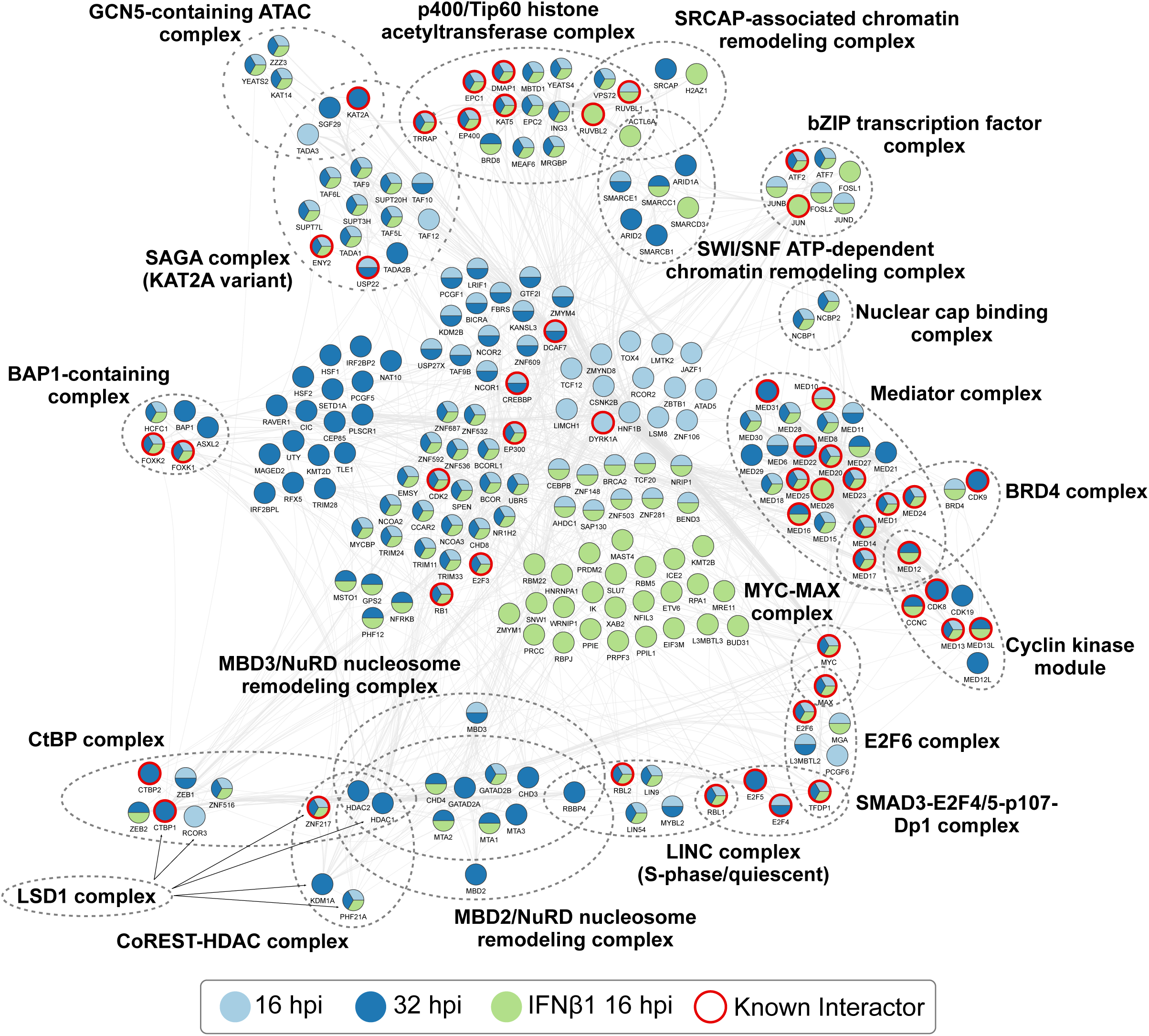
Protein complex analysis of E1A interactors. Integrated E1A PPI network organized based on protein complex membership. High-confidence E1A PPIs were analyzed with the Cytoscape ClueGO plug-in to identify statistically enriched protein complexes annotated in the CORUM or Complex Portal databases. Protein nodes are colored according to time point or IFNβ1 treatment and known E1A interactors are indicated with a red border. PPI interaction evidence from the IntAct database was used to construct the protein interaction network.

We identified nearly all subunits of the p400 complex (**Fig. S4A**), a 17-subunit assembly in which p400 scaffolds several modular components, as recently defined by cryo-EM^58–60^. This included interactions with BRD8 and MRGBP, two subunits of the Tintin module that have not previously been shown to interact with E1A. Notably, the Tintin module has failed to co-purify with the p400 complex in recent cryo-EM structures, presumably reflecting a transient association. Mediator was similarly well represented. We identified 24 of 26 subunits of the core Mediator complex^61^, along with the cyclin kinase module and the Brd4 complex^62^, and observed putative novel interactions with CDK19 and MED12L, which are subunit paralogs of the cyclin kinase module^63^. Mediator is a well-established E1A target, essential for E1A-mediated transactivation, with the interaction previously mapped to E1A conserved region 3^44,45^. Considering our approach recovers this established complex to near completion alongside the p400 complex, whose Tintin module appears transient, this demonstrates our strategy captures both stable and transient components of large host complexes with comparable sensitivity.

### Proximity Labeling Resolves Transient Interactions

To directly test whether our approach could resolve interactions less amenable to conventional methods, we compared detection of the previously unreported interaction of E1A with BRD8 by immunoprecipitation and by BioID in an infection context (**Fig. S4B**). We were able to detect the E1A-BRD8 interaction during infection by BioID and western blotting using our FmT-E1A virus, but not by direct E1A immunoprecipitation. Performing a reciprocal immunoprecipitation of BRD8 confirmed the interaction (**Fig. S4C**), validating BRD8 as a genuine E1A interactor that would have been missed by E1A immunoprecipitation alone. This interaction appeared preferential to the 13S E1A isoform, which is substantially less abundant than 12S E1A at 24 hpi, suggesting isoform-specific specificity for the Tintin module. Our approach similarly recovered Tip60/Kat5, which was previously shown to interact with E1A, albeit only in transient overexpression studies^64^, further demonstrating sensitivity for interactions that are difficult to capture using standard approaches in an infection context.

We extended this validation to the nuclear cap-binding complex, composed of NCBP1 and NCBP2, which is a key regulator of RNA processing^65,66^. We tracked this interaction by BioID and western blotting across a time course of infection and found clear enrichment relative to the FmT-2A control at every time point tested (**Fig. S4D**). Notably, NCBP3, a third, less well-characterized nuclear cap-binding paralog^67^, was quantified in our dataset but did not meet our high-confidence threshold, indicating that E1A preferentially engages NCBP1/2 over NCBP3. Together, these data show that our strategy can identify novel interactors, resolve both weak/transient interactions missed by immunoprecipitation, and help define specificity between closely related host proteins.

### The p400 Complex is the Dominant E1A Target During the Immediate-Early Phase of Infection

E1A-host PPIs characterized to date have largely been studied toward the end of the early phase or into the late phase. Like most immediate-early viral proteins, the repertoire of E1A interactions formed within the essential first hours of infection, when abundance is at its lowest, has not been investigated. To resolve this window, we provided exogenous biotin immediately following infection (1 hpi), preceding E1A gene expression^68,69^, and allowed labeling to continue until 8 hpi (**Fig. 4A**). This strategy leverages the catalytic nature of biotinylation, such that biotinylated E1A interactors will exist in stoichiometric excess relative to E1A at the culmination of our labeling window. In essence, this will “amplify” the PPI signal from a bait present at minimal abundance via sequential transient interactions with E1A targets.

**Figure 4.**
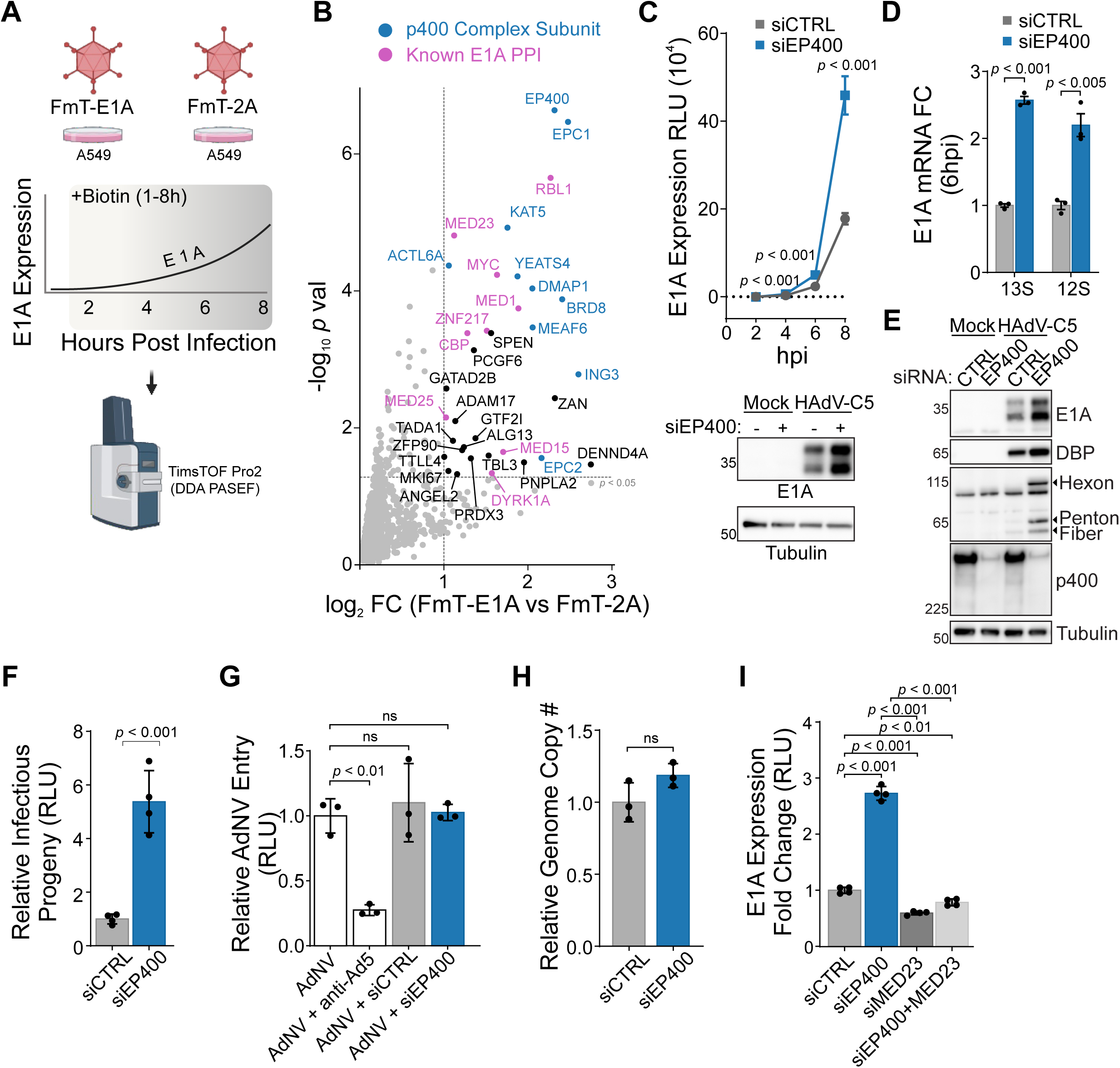
p400 is the dominant target of E1A during immediate early infection and restricts E1A expression. **A)** Immediate early phase E1A interactome experiment design schematic. A549 cells were infected with FmT-E1A or -2A virus and biotinylation was carried out from 1 hpi to 8 hpi when cells were harvested for streptavidin enrichment and identification by mass spectrometry. **B)** Volcano plot of high confidence E1A interactors (Log2 FC ≥ 1, *p* val ≤ 0.05) identified during the immediate early phase of infection. Subunits of the p400 complex are labeled blue and additional known E1A interactors are labeled in magenta. Proteins labeled in black represent putative novel E1A interactors. **C)** Time course of E1A protein expression, measured by nanoluciferase activity, in cells infected with AdNTE reporter virus. Cells were transfected with siRNA control (siCTRL) or siEP400 for 24 hours prior to infection. A corresponding western blot from 8 hpi is displayed below. Multiple *t* test with Holm Sidak adjustment, n = 6. **D)** RT-qPCR quantification of E1A mRNA (13S and 12S isoforms) at 6 hpi in A549 cells knocked down for p400 and infected with HAdV-C5. Fold change (FC) is relative to siCTRL. Multiple *t* test with Holm Sidak adjustment, n =3. **E)** Viral protein expression at 18 hpi in A549 cells infected with HAdV-C5 following 24 hours of p400 knockdown. **F)** Infectious progeny production following 24 hours of p400 knockdown using AdNTE reporter virus. *t* test, n = 4. **G)** Relative quantification of viral entry at 20 minutes post-infection following 24 hours of p400 knockdown. Entry was measured by infecting A549 cells with the AdNV reporter virus which uses bioluminescent virions (nanoluciferase) to quantify virus uncoating upon entry. One-way ANOVA with Dunnett’s correction, n =3. **H)** Relative quantification of nuclear HAdV-C5 genomes at 4 hpi determined by qPCR following 24 hours of p400 knockdown. *t* test, n =3. **I)** E1A protein expression at 6 hpi following p400 and/or MED23 knockdown. A549 cells were infected with AdNTE reporter virus after 24 hours of siRNA knockdown and E1A expression was measured by nanoluciferase activity. One-way ANOVA with Tukey’s correction, n =4.

Analysis of high-confidence PPIs from this immediate-early window identified multiple subunits of the p400 complex as the top differentially enriched interactors by both fold-change and significance (**Fig. 4B**, Supplemental Data 4). We recovered all four subunits of the HAT module (EPC1, KAT5, MEAF6, and ING3) as well as subunits of the ARP and Tintin modules. TRRAP, while quantified in all samples, did not meet our high-confidence threshold and may reflect a temporally distinct p400 complex composition specific to this stage of infection. This finding was unexpected considering p300/CBP, Mediator, and Rb are the classical E1A interactors associated with viral early gene expression and cell-cycle control^70^. Notably, while we identify these proteins as high confidence PPIs, they were of lower fold-change compared to p400 at the earliest times during infection.

In addition to p400, we identified several transcriptional regulators at these earliest timepoints that have not previously been linked to E1A, including the transcriptional repressors SPEN and PCGF6^71,72^. We also identify GTF2I, an established HAdV restriction factor targeted by E4orf3 for degradation later in infection^73,74^. GTF2I was also recently shown to bind the adenovirus genome directly and restrict infection independent of the DNA-damage or interferon responses^75^. Identification of GTF2I at this stage supports GTF2I acts as an intrinsic antiviral factor. We recovered known E1A interactors, including RBL1, MYC, CBP, ZNF217, and the Mediator subunits MED1, MED15, MED23, and MED25. Interestingly, MED15, MED23, and MED25 belong to the Mediator tail module, which docks transcription factors onto the core Mediator complex^76,77^. MED23 specifically is the primary contact point between E1A conserved region 3 and Mediator^45^; here, the selective biotinylation of tail-module subunits at this stage directly supports the known topography of E1A-Mediator engagement. Together, these data reveal that the immediate-early E1A interactome is functionally diverse, spanning chromatin remodelers (p400 complex), transcriptional repressors (PCGF6 and SPEN), activators (Mediator complex), and a known host restriction factor (GTF2I).

### p400 Restricts E1A Expression Independent of Viral Entry, Genome Delivery, or Global Proteome Effects

Given that p400 emerged as E1A’s dominant immediate-early target and an essential role of E1A is regulating viral early gene expression (including E1A itself), we hypothesized this interaction contributes to transcriptional regulation. We performed siRNA-mediated knockdown of p400 in A549 cells (**Fig. S5A**; siEP400#1 used throughout) and infected cells with a nanoluciferase-P2A-E1A reporter virus (AdNTE)^78^ to track E1A expression across the immediate-early phase (**Fig. 4C**). Contrary to our hypothesis, knockdown of p400 resulted in a greater than two-fold increase in E1A protein expression, which was detectable as early as 4 hpi (**Fig. S5B**). We also observed a similar increase in E1A protein following AdNTE or wild-type HAdV-C5 infection of primary human bronchial epithelial cells (HBEC) at 6 and 18 hpi, respectively (**Fig. S5C and S5D**). Furthermore, p400 knockdown in HEK293 cells, which constitutively express E1A from an integrated HAdV-C5 left-end fragment, similarly increased E1A protein but to a lesser extent (**Fig. S5E**). RT-qPCR at 6 hpi similarly showed E1A mRNA increased 2–3-fold following p400 knockdown (**Fig. 4D**). By 18 hpi, elevated E1A was accompanied by increased expression of the early protein DBP, which is transcriptionally regulated by E1A, as well as late capsid proteins (Hexon, Penton, and Fiber) (**Fig. 4E**). These observations also coincided with a nearly 6-fold increase in infectious progeny production as early as 18 hpi (**Fig. 4F**), suggesting p400 knockdown accelerates progression through the replication cycle. Together, these data establish that p400 restricts E1A expression during the immediate-early phase, with functional consequences that propagate through the full replication cycle.

Since p400 knockdown might increase E1A expression through indirect cellular effects rather than a direct restrictive function, we next tested several alternative explanations. Viral entry, measured using a bioluminescent-virion reporter virus (AdNV)^78^, and nuclear delivery of viral genomes at 2 hpi were both unaffected by p400 knockdown (**Fig. 4G and 4H**), ruling out an entry- or nuclear trafficking-based mechanism. Cell-cycle distribution in uninfected cells was unchanged 24 hours after p400 knockdown (**Fig. S6A**), and whole-cell proteomics of uninfected cells identified only 12 differentially abundant proteins, none of which are known E1A targets (**Fig. S6B**, Supplemental Data 5). These data imply that global cellular effects of p400 depletion are minimal within this timeframe. The whole-cell proteomics quantified 15 of the 17 p400 complex subunits, all of which remained stable upon p400 knockdown (**Fig. S6C**), suggesting that the restrictive phenotype we observe reflects loss of p400 itself, or its capacity to scaffold an intact complex, rather than degradation of orphaned subunits^79^. Collectively, these controls indicate that p400 restricts E1A expression directly.

The immediate-early E1A interactome is composed of very large proteins and complexes. These include CBP (300 kDa), and the megadalton-scale p400 (1.7 MDa) and Mediator (1.2–2 MDa) complexes (**Fig. 4B**). The CBP-binding region of E1A is contiguous with its p400-binding region, and E1A may also engage CBP at a secondary site that overlaps with the Mediator-binding site within conserved region 3^37^. Given this overlap, it is unlikely that a single E1A molecule binds all three simultaneously, and it is more probable that these represent mutually exclusive interactions competing for a limited pool of E1A (*i.e.,* PPI epistasis). Under this model, when E1A abundance is minimal during the immediate-early phase, the interactions that dominate should be those most kinetically or stoichiometrically favored. Because Mediator subunits were enriched at a lower fold-change than p400 in our immediate-early dataset, we hypothesized that loss of p400 would shift this competition toward E1A-Mediator engagement, accounting for the increase in E1A expression upon p400 knockdown. Consistent with this model, combined knockdown of p400 and MED23, the primary contact point between E1A and the Mediator complex, abolished the increase in E1A expression otherwise seen with p400 knockdown alone (**Fig. 4I and S7A**). Similarly, targeting CBP directly with the degrader dCBP-1^80^ decreased E1A expression when p400 has been knocked down (**Fig. S7B and S7C**). These experiments, together with our immediate-early BioID data, indicate that E1A expression at this stage is governed by a competitive, stoichiometry-driven network of interactions rather than by p400 acting in isolation.

### p400 Associates with Incoming Adenovirus Genomes

Sensitive measurements of E1A mRNA and protein have shown that E1A expression becomes detectable above background only after 2 hpi^69,78^. Considering viral genomes are delivered to the nucleus by 1 hpi^81^, there is a considerable gap between nuclear genome delivery and E1A gene expression. Since p400 knockdown increased E1A protein as early as 4 hpi, we hypothesized that p400 might act directly on incoming nuclear genomes, prior to viral early gene transcription. To test this, we performed p400 chromatin immunoprecipitation followed by quantitative PCR (ChIP-qPCR) targeting viral genomic regions, including the E1A promoter and enhancer, the internally located tripartite leader (TPL), and the Fiber open reading frame, which lies at the right end of the HAdV genome. At 2 hpi, p400 was significantly enriched at the E1A enhancer and promoter relative to both GAPDH and Fiber (**Fig. 5A**). We also detected E1A at the TPL, indicating that p400 associates broadly with the incoming genome rather than exclusively at the E1A promoter/enhancer. To determine if p400 might non-specifically restrict expression of other viral early genes independent of E1A, we knocked down p400 and infected cells with WT or an E1A deleted virus, *dl*312 (**Fig. S8A**). During *dl312* infection, some DBP protein expression was observed, and this was similarly increased following p400 knockdown. We do not see full rescue of DBP expression; however, this is not surprising because E1A and E4orf6/7 are required for full activation of DBP^82^, neither of which are expressed during *dl312* infection. Nevertheless, these data suggest p400 restriction is not limited to the E1A enhancer/promoter. We also performed ChIP-qPCR at 1 hpi and similarly observed p400 associated with the viral genome (**Fig. S8B**), providing greater confidence p400 can associate with incoming viral genomes independent of E1A expression.

**Figure 5.**
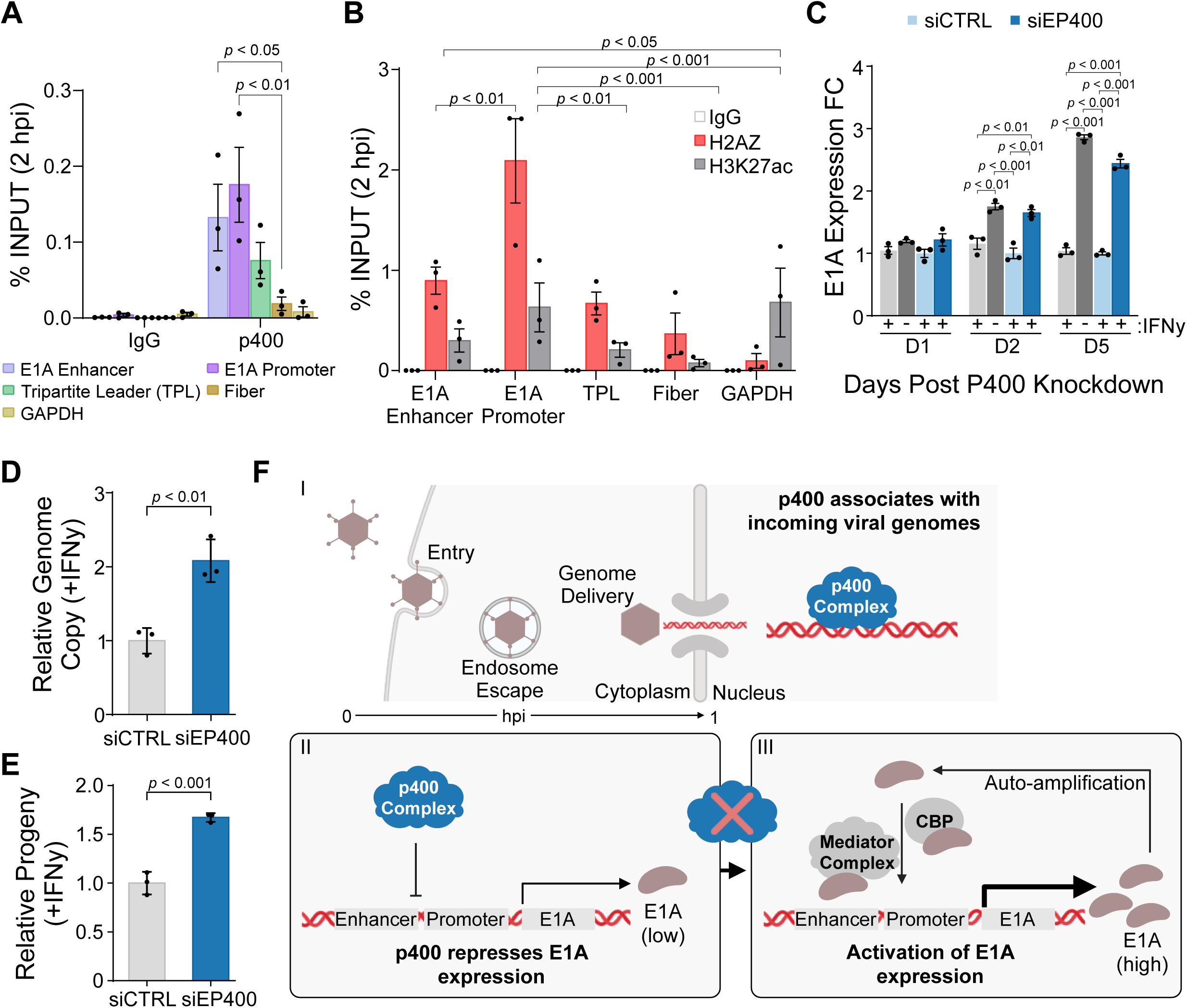
p400 is associated with incoming adenovirus genomes. **A)** p400 chromatin immunoprecipitation (ChIP) qPCR at 2 hpi in A549 cells infected with HAdV-C5. Fiber open reading frame was used as the control for statistical comparisons. Only significant tests are shown (p < 0.05). One-way ANOVA with Fisher LSD, n =3. **B)** H2AZ ChIP-qPCR at 2 hpi in A549 cells infected with HAdV-C5. GAPDH was used as a negative control for H2AZ and a positive control H3K27ac. Statistical comparisons were made for H2AZ only. Only significant tests are shown (p < 0.05). One-way ANOVA with Fisher LSD, n =3. **C – E)** Persistent HAdV infection model in HDF-TERT cells infected with AdNTE in the presence of IFNγ. At 30 days post-infection cells were transfected with siEP400. E1A expression was assessed up to 5 days post p400 knockdown (C). E1A expression fold change (FC) is normalized to siCTRL (light blue). Grey bars represent untransfected cells. Genome replication (D) and infectious progeny production (E) were quantified at 7 days post p400 knockdown. Panel C; One-way ANOVA with Tukey’s correction, n = 3. Panel D and E; *t* test, n = 3. **F)** Schematic representing influence of p400 knockdown on E1A expression. P400 associates with incoming viral genomes and represses E1A expression. Knockdown of p400 increases E1A expression, which enhances E1A-mediated regulation of the E1A promoter via interactions with Mediator and CBP.

As an independent test of this association, we performed ChIP-qPCR for the histone variant H2AZ, which regulates transcription and can be deposited by the p400 complex^83^. Using GAPDH regions validated as H2AZ-negative and H3K27ac-positive as controls^84^, we observed H2AZ localization at multiple HAdV genomic regions at both 1 and 2 hpi (**Fig. 5B and S8C**). Together, these data show that p400 engages the adenovirus genome upon nuclear entry and prior to the onset of viral gene expression, positioning it as a poised restriction mechanism rather than an E1A recruited factor.

### p400 Restricts E1A Expression to Maintain Persistent Adenovirus Infection

The E1A enhancer/promoter is known to be suppressed by interferon signaling, a repression that is critical for HAdV to establish persistent infection. Furthermore, HAdV replication can be rescued in the presence of IFNγ by E1A overexpression^28,85^, indicating that E1A repression is itself a determinant of persistence and the possibility that the E1A p400 interaction antagonizes this process to promote acute infection. Because p400 restricts E1A expression during acute infection, we asked whether it similarly contributes to persistence. Using a previously established persistence model^85^, we infected TERT-immortalized human dermal fibroblasts (HDF) with the AdNTE reporter virus and maintained cultures in IFNγ for 30 days before transfecting with siEP400 or control siRNA. We validated the persistence model independently by removing IFNγ from non-siRNA transfected cultures at 30 days post-infection, which produced a 3-fold increase in viral genome copy number (**Fig. S8D**), confirming that persistence in this system is actively maintained by ongoing interferon signaling.

Following p400 knockdown in the persistent infection model, we observed a time-dependent increase in E1A expression alongside increased viral genome replication and infectious progeny production (**Fig. 5C–E**). These data indicate p400 continues to restrict E1A and constrain viral output during persistence. p400 knockdown similarly rescued E1A protein expression following acute IFNβ treatment of A549 cells (**Fig. S8E**), showing that this restrictive function operates across both the acute and persistent states. In summary, these data identify p400 as an early host restriction factor and a regulator of HAdV persistence, acting through repression of E1A expression. We propose a model in which p400 associates with incoming viral genomes immediately upon nuclear entry and attenuates E1A expression (**Fig. 5F**). Loss of p400 permits accelerated E1A auto-activation through its downstream targets (*e.g.,* Mediator and CBP), and in the context of interferon-driven persistence, this p400-mediated restraint helps maintain the limited E1A expression required to sustain persistent infection.

## Discussion

The immediate-early phase disproportionately determines the outcome of infection and has remained largely inaccessible to virus-host PPI mapping. During this phase, incoming viral genomes are being surveilled by the host, typically involving chromatinization and engagement of transcription regulators^86–88^. Critically, these events occur before viral protein abundance is sufficient for standard interactome methods to adequately resolve them. Identifying the repertoire of host proteins that engages viral DNA and the viral proteins that regulate these earliest hours is central not only to understanding how a productive infection is established, but to defining intrinsic host defenses more broadly.

Our approach offers several advances over existing strategies for mapping virus-host PPIs in an infection context. The paired isogenic P2A control virus provides a background that is matched simultaneously for bait abundance and infection-driven remodeling of the host cell proteome, addressing a confounding variable that mock infection, IgG, or ligase-only controls poorly resolve. By leveraging the catalytic properties of a genetically encoded biotin ligase we could focus our investigation onto the immediate-early phase of infection when bait abundance is lowest. Importantly, because labeling occurs during authentic infection, the interactions we capture reflect endogenous stoichiometry and native subcellular context. Using this approach, we investigated the immediate-early E1A interactome and discovered p400 was the dominant E1A target at this stage. More broadly, our data highlight the remarkable ability of a small viral protein to concurrently dysregulate cell cycle (Rb), initiate viral gene expression (Mediator), and antagonize host restriction factors (p400), a feat typically requiring multiple viral proteins.

More than two decades ago p400 was discovered via PPI studies with E1A^38^. Subsequent work has linked this interaction to E1A-driven cellular transformation and apoptosis^89,90^, stabilization of Myc^40^, and, most recently, activation of Alu retrotransposon transcription^91^. Yet despite two decades of study, what this interaction accomplishes for the virus during infection has remained undefined. Our data suggests a different role for this interaction, where p400 first functions as a restriction factor that E1A must overcome. In the earliest stages of infection, p400 attenuates E1A expression, in an attempt to abort infection given the key role of E1A in activating early gene expression. The limited amount of E1A which is expressed and translated at this point appears to preferentially and directly antagonize p400-mediated repression to establish a successful acute lytic infection that yields high levels of viral progeny. In the presence of interferon, this balance is shifted and the virus enters a long-term persistence state characterized by a low level of virus replication and little cell death.

Concurrently to the targeting of p400, E1A molecules can separately engage factors like Mediator and CBP/p300 to drive auto-activation of the E1A promoter. This model is consistent with our data showing that p400 is the dominant E1A target and E1A-Mediator engagement, rather than loss of p400 itself, accounts for the increase in E1A expression following p400 knockdown.

Host factors that contribute to the temporal gap between HAdV nuclear genome delivery and viral early gene expression remain poorly characterized. Host intrinsic defenses targeting incoming viral genomes have been documented for several DNA viruses and this is best described with herpesviruses and PML nuclear bodies^92,93^, which restrict the viral genome. Incoming HAdV genomes were initially suspected to associate with PML nuclear bodies^94^; however, it was later shown incoming Ad genomes neither recruit PML nuclear body components nor colocalize with them^95^. Our data agrees with a PML-independent model and identifies p400, along with other putative host proteins, as a novel HAdV restriction factor. Whether this restriction is related to histone deposition (*e.g.,* H2AZ) or other epigenetic functions carried out by the p400 complex (*e.g.,* acetylation) is unknown.

There are several hypothetical mechanisms by which p400 might be recruited to the viral genome. Notably, incoming genomes are packaged with a viral histone, pVII^96^, which may be recognized by p400. However, the E1A promoter region was recently shown to be the least enriched for pVII on incoming viral genomes^68^. Why genomes are packaged this way is intriguing, and it’s possible that limiting viral histone deposition at the E1A promoter helps facilitate transcription in subsequent rounds of infection. However, in such a scenario, this nucleosome devoid region could be recognized by p400 as a protective “gap filling” measure, like what has been observed with HIRA and other DNA viruses, albeit in a PML-dependent manner^97^.

The p400 complex is emerging as an important regulator for viral infection and has been shown to be targeted by HIV-1 Tat^98^, HPV18 E2^99^, HTLV-1 p30II^100^, and MCPyV small T antigen^101^, among others^102,103^. While some of these models have shown p400, or at least one of the subunits (*e.g.,* Kat5/Tip60), can associate with viral genomes, these studies have primarily investigated latent or late phase time points of infection. Furthermore, for Tat and E2, transcriptional repression is viral protein dependent. In contrast to these other viral systems, our data demonstrates p400 can associate with incoming viral genomes and that this precedes early gene expression. In addition to the E1A enhancer/promoter, p400 and H2AZ also occupied the TPL and Fiber genomic regions, which suggests p400 may act broadly upon the viral genome.

In summary, we provide a pathogen-driven proximity proteomics framework that resolves virus-host PPI networks with spatial and temporal precision previously inaccessible to existing approaches. Using our approach, we identified p400 as a poised host restriction factor acting directly on the incoming viral genome. More broadly, roughly one-third of the high-confidence interactors identified in this study are not annotated as targeted by any other virus family, while the remaining two-thirds are shared across multiple viral taxa, suggesting this dataset has additional value for virus-host interactions beyond adenovirus. This strategy can be extended to other viral systems, and the P2A-based normalization strategy we describe offers a solution for controlling bait abundance and background proteome changes associated with infection, which are central to interactome studies in a dynamic system.

## Materials and Methods

### Reagents and materials

See Reagents Table for a complete list of reagents and materials used in this study.

### Tissue culture, infections, and siRNA transfection

A549, HDF-TERT, and HEK 293 cells were grown in DMEM supplemented with 10% fetal bovine serum and 1% pen/strep at 37°C with 5% CO2. HBEC-KT3 cells were grown in Airway Epithelial Cell Basal Medium supplemented with Bronchial Epithelial Cell Growth Kit and 1% v/v Pen/Strep. All viral infections were carried out at an MOI of 10 in a minimal volume of serum free media for 1 hour. All cell lines were routinely tested for mycoplasma contamination. siRNA knockdown was performed by reverse transfection using Lipofectamine RNAiMax. siRNA used in this study are available in the Primers and siRNA Table.

### HAdV-C5 persistence model

HDF-TERT cells were treated with 1000 U/mL IFNy for 24 hours and then infected with the AdNTE reporter virus (MOI 10). Cells were maintained in DMEM (10% FBS, 1% Pen/Strep) for 30 days without passaging. Every three days an additional 1000 U/mL IFNy was added. At 30 days post infection cells were split and reverse transfected with siRNA.

### Plasmids and viruses

Wild-type HAdV-C5 was obtained from the ATCC. AdNTE and AdNV reporter viruses were acquired from Dr. Andrew Mehle^78^. AdNTE virus expresses E1A as a nanoluciferase-P2A fusion and AdNV expresses pV, which is incorporated into packaged virions, as a nanoluciferase-pV fusion. Construction of FmT-E1A and FmT-2A-E1A viruses was performed following the Adenobuilder protocol for HAdVC-5^104^. In brief, FmT and FmT-P2A coding sequences were first cloned into pAd5-B1 plasmid by Gibson assembly and validated by Sanger sequencing. Sequenced verified pAd5-B1 plasmids are available in supplementary files. FmT was cloned with a 3xFLAG and SGGSGG linker/spacer between FmT and E1A. Plasmids for pAd5-B1, -B2, -B3/4, -B5 and -B6/7 were digested with BstBI, Gibson assembled using NEBuilder HiFi DNA Assembly Master Mix and transfected into two million HEK 293 cells by electroporation. At 5 days post transfection cells and supernatant were freeze-thawed three times and a second round of infection was performed on HEK 293 cells to amplify rescued virus. Rescued virus was subsequently plaque purified on HEK 293 cells and Sanger sequenced to confirm miniTurbo insertion. Validated viruses were propagated and purified on cesium-chloride gradient.

### Confocal immunofluorescence microscopy

A549 cells were seeded onto glass coverslips and infected 24 hours later. Cells were washed twice in PBS, fixed in 4% paraformaldehyde solution for 10 minutes at room temperature and washed 3 times in PBS. For infected cells that were pre-extracted, cells were incubated on ice for one minute with CSK buffer prior to fixation. Cells were permeabilized in IF permeabilization buffer at room temperature for 10 minutes, washed in TBS+0.1% Tween-20 (TBST), and blocked in 3% BSA+TBST for 1 hour. Coverslips were incubated overnight at 4°C with primary antibodies, washed 3x in PBS, and incubated with AlexaFluor secondary antibodies in 3% BSA+TBST (1:1000 dilution) and DAPI (0.1 µg/mL) for 1 hour at room temperature. Coverslips were washed 3x in PBS and once with water and mounted with ProLong Gold Antifade Mountant. Confocal immunofluorescence was performed using a Zeiss LSM 980 confocal equipped with a 63x/1.4 NA oil immersion objective.

### Immunoprecipitation

Six Million A549 cells in 10cm dishes were infected with wild type HAdV-C5. At 24 hpi, cells were scraped and cell pellets were lysed in 500uL NP-40 lysis buffer supplemented with protease inhibitor cocktail and Benzonase (1:1000), immediately freeze-thawed in liquid nitrogen, and incubated for 30 minutes at 4°C with end-over-end rotation. Lysed samples were centrifuged to remove debris, and supernatant was transferred to fresh 1.5 mL Eppendorf tubes containing antibody-bead complexes. Antibody-bead complexes were prepared by washing protein A or G dynabeads in NP-40 lysis buffer and incubating with antibody for 1 hour at room temperature. Bead-antibody complexes were mixed using 15uL of Dynabead per 1 µg antibody. Following binding, beads were washed once in NP-40 lysis buffer. Immunoprecipitations were carried out at 4°C for 2 hours with end-over-end rotation and washed three times in NP-40 lysis buffer. Beads were resuspended in 2x LDS sample buffer containing BME and boiled for 10 minutes. Samples were resolved on a 4-12% NuPAGE Bis-Tris protein gel and transferred to PVDF for western blot.

### BioID sample preparation and mass spectrometry

Twelve million A549 cells in 15cm dishes were infected with FmT-E1A or FmT-2A-E1A virus at MOI 10 pfu/cell. Two dishes per biological replicate (n=3) were seeded for each condition and pooled. For cells treated with IFNβ1, 1 U/mL was added to media 16 hours prior to infection and 1 U/mL was maintained over the course of infection. Biotin was spiked into media containing infected cells at a final concentration of 50 µM for two hours. For immediate-early phase E1A interactome (1-8 hpi), biotin was added immediately following infection (1 hpi) at a 50 µM final concentration until time of harvest at 8 hpi. Cells were washed twice in PBS, scraped, snap frozen in liquid nitrogen, and their weight was recorded. Lysis was performed by adding RIPA lysis buffer supplemented with protease inhibitor cocktail and Benzonase (1:1000) at a 6:1 ratio to cell pellet mass (e.g., 600 µL/100 mg). Samples were incubated on ice for 30 minutes with periodic vortexing, centrifuged to clear cell debris, and transferred to fresh LoBind Eppendorf tubes containing 30 µL streptavidin sepharose beads which have been pre-washed in RIPA lysis buffer. Streptavidin pulldown was performed for 4 hours at 4°C with end-over-end rotation.

Streptavidin enriched samples were washed twice in RIPA buffer, transferred to fresh LoBind Eppendorf tubes and washed an additional 3 times in 50 mM ABC. Beads were resuspend in 100 µL 50 mM ABC containing 1 µg trypsin to carry out on-bead digestion overnight at 37°C on a thermomixer. The following day an additional 0.5 µg trypsin was added to samples and incubated for an additional 3 hours. Sepharose beads were centrifuged at 250g for 1 minute and 100 µL of supernatant containing peptides was transferred to fresh LoBind Eppendorf tubes. Sepharose beads were washed twice in 100 µL HPLC water and pooled. Formic acid was added to peptides (5% final) and peptides were dried by vacuum centrifugation.

BioID samples from corresponding to 16hpi, 32hpi, and IFNB treatment were acquired on a TripleTOF 6600 (Sciex) in both DDA and DIA. For data-dependent acquisition (DDA) LC-MS/MS, affinity purified and digested peptides were analyzed using a nano-HPLC (High-performance liquid chromatography) coupled to MS. One-quarter of the sample was used. Nano-spray emitters were generated from fused silica capillary tubing, with 100µm internal diameter, 365µm outer diameter and 5-8µm tip opening, using a laser puller (Sutter Instrument Co., model P-2000, with parameters set as heat: 280, FIL = 0, VEL = 18, DEL = 2000). Nano-spray emitters were packed with C18 reversed-phase material (Reprosil-Pur 120 C18-AQ, 3µm) resuspended in methanol using a pressure injection cell. Sample in 5% formic acid was directly loaded at 800nl/min for 20min onto a 100µmx15cm nano-spray emitter. Peptides were eluted from the column with an acetonitrile gradient generated by an Eksigent ekspert™ nanoLC 425, and analyzed on a TripleTOF™ 6600 instrument (AB SCIEX, Concord, Ontario, Canada). The gradient was delivered at 400nl/min from 2% acetonitrile with 0.1% formic acid to 35% acetonitrile with 0.1% formic acid using a linear gradient of 90 min. This was followed by a 15 min wash with 80% acetonitrile with 0.1% formic acid, and equilibration for another 15min to 2% acetonitrile with 0.1% formic acid. The total DDA protocol is 135min. The first DDA scan had an accumulation time of 250ms within a mass range of 400-1800Da. This was followed by 10 MS/MS scans of the top 10 peptides identified in the first DDA scan, with accumulation time of 100 ms for each MS/MS scan. Each candidate ion was required to have a charge state from 2-5 and a minimum threshold of 300 counts per second, isolated using a window of 50mDa. Previously analyzed candidate ions were dynamically excluded for 7 seconds. For data-independent acquisition (DIA) LC-MS/MS, affinity purified and digested peptides were analyzed using a nano-HPLC (High-performance liquid chromatography) coupled to MS. One-quarter of the sample was used. Nano-spray emitters were generated from fused silica capillary tubing, with 100µm internal diameter, 365µm outer diameter and 5-8µm tip opening, using a laser puller (Sutter Instrument Co., model P-2000, with parameters set as heat: 280, FIL = 0, VEL = 18, DEL = 2000). Nano-spray emitters were packed with C18 reversed-phase material (Reprosil-Pur 120 C18-AQ, 3µm) resuspended in methanol using a pressure injection cell. Sample in 5% formic acid was directly loaded at 800nl/min for 20min onto a 100µmx15cm nano-spray emitter. Peptides were eluted from the column with an acetonitrile gradient generated by an Eksigent ekspert™ nanoLC 425, and analyzed on a TripleTOF™ 6600 instrument (AB SCIEX, Concord, Ontario, Canada). The gradient was delivered at 400nl/min from 2% acetonitrile with 0.1% formic acid to 35% acetonitrile with 0.1% formic acid using a linear gradient of 90 min. This was followed by a 15 min wash with 80% acetonitrile with 0.1% formic acid, and equilibration for another 15min to 2% acetonitrile with 0.1% formic acid. The total DIA protocol is 135min. The first DIA scan had an accumulation time of 250ms within a mass range of 400-1800Da. This was followed by 54 MS/MS scans with differing mass windows, with an accumulation time of 65ms per scan. DDA and DIA data were searched with MSFragger-DIA111 (default settings) using the human proteome (UP000005640), containing both SWISS-PROT and TrEMBL entries, and the human adenovirus 5 proteome (UP000004992). Oxidation of methionine and N-term acetylation were set as variable modifications.

Immediate-early phase BioID samples, corresponding to 1-8hpi, were acquired on a Bruker TimsTOF Pro 2 in DDA-PASEF. One-sixteenth of digested peptides were analyzed using a nano-HPLC coupled to MS. The sample was loaded onto Evotip Pure per manufacturer instructions. Peptides were eluted from the Performance column (cat#: EV-1109, 8cmx150µm with 1.5µm beads), with the 60SPD pre-formed acetonitrile gradient generated by an Evosep One system. The Evosep was coupled to timsTOF Pro 2 using a 20um diameter emitter tip. The column toaster was set to 40C. The total DDA protocol is 22 minutes. The MS1 scan had a mass range of 100-1700Da in PASEF mode. TIMS settings were accumulation and ramp time of 100ms (with 4 PASEF ramps and active exclusion at 0.4min), and within the mobility range (1/K0) of 0.85 to 1.3V·s/cm2. This was at a cycle time of 0.53s. The target intensity was set to 17,500 and intensity threshold set to 1750. 1+ ions are excluded from fragmentation using a polygonal filter. The auto calibration was off. Mass spectra were searched with MSFragger 4.1 within the ProHits LIMS. FASTA database is from UP000005640 human Uniprot, no isoforms, and HAdV-C5 proteome. Acetylated protein N-term and oxidated methionine were set as a variable modifications. Precursor mass tolerance was set to 20ppm on either side. Fragment mass tolerance was set to 20ppm. Enzymatic cleavage was set to trypsin with 2 missed cleavages. MSBooster and Percolator were turned on. Percolator required a minimum probability of 0.5 and did not remove redundant peptides. The target-decoy competition method was used to assign q-values and PEPs. For ProteinProphet, the maximum peptide mass difference was set to 30ppm. When generating the final report, the protein FDR filter was set to 0.01. FDR was estimated by using both filtered PSM and protein lists. Razor peptides were used for protein FDR scoring. All other parameters were default.

Protein groups were filtered to remove contaminants and those with high missing values. Protein intensities were log2 transformed, vsn normalized, and imputed using minimum probably distribution. Differential protein abundance was performed with Limma. Proteins with a log2 fold-change ≥ 1 compared to control (FmT-2A) and p.val, or p.adj, ≤ 0.05 were considered high confidence protein interactors.

### P400 knockdown whole cell proteomics

Four-hundred thousand A549 cells were reverse transfected with a control siRNA or siEP400 #1. At 24 hours post transfection A549 cells were harvested by trypsinization and cell pellets were lysed in 30 uL 5% SDS. Samples were sonicated in a Biorupter Pico (10x 30 seconds ON/OFF). Protein concentration was determined by BCA assay using the Pierce Rapid Gold BCA kit. Further processing of samples was performed using an S-TRAP micro column following the manufacturer’s instructions and dried by vacuum centrifugation.

Samples were acquired on a Bruker TimsTOF Pro 2 in DIA-PASEF. For data-independent acquisition (DIA) LC-MS/MS, 250ng of digested peptides were analyzed using a nano-HPLC coupled to MS. 250ng of the sample was loaded onto Evotip Pure per manufacturer instructions. Peptides were eluted from the column (cat#: EV-1137, 15cmx150µm with 1.5µm beads) with the 30SPD pre-formed acetonitrile gradient generated by an Evosep One system. The column toaster was set to 40C. The total DIA protocol is 44 minutes. The MS1 scan had a mass range of 100-1700Da in dia-PASEF. TIMS settings were accumulation and ramp time of 100ms, and within the mobility range (1/K0) of 0.6 to 1.6V·s/cm2. Cycle time 2s. For MS2, 1 mobility window with 2 ramps was used for 32 mass windows, 29.8 Da wide with 5Da mass overlap. The mobility range was from 0.61/K0 to 1.451/K0. This was at a duty cycle of 100% and a ramp rate of 9.52Hz. 1+ ions are excluded from fragmentation using a polygonal filter. The auto calibration was off.

Spectronaut v20 directDIA+ BGS workflow was used to search the data with the Spectronaut generated human spectral library (Human_PDB_2023). Parameters for the search were default, normalization was on. Differential abundance testing was performed with Limma.

### Flow cytometry

Propidium iodide staining was used to quantitate cell cycle stages following siRNA transfection. Cells were transfected with siRNAs for 24 hours before fixing in 95% ice cold ethanol. Ethanol was added dropwise while vortexing resuspended cells to a final concentration of 70% ethanol. Cells were fixed at 4°C for 30 minutes on ice. Cells were then washed twice with PBS to remove any remaining ethanol prior to resuspending in RNase/Staining buffer. Next, propidium iodide (1mg/mL) was added 1:50 to the cells (final concentration 20 ug/mL) and stained for 15 minutes at 37°C with rotation at 600 rpm. Cells were then filtered through a 5 mL tube with a 12x75mm cell strainer cap and sorted within 48 hours post propidium iodide staining. Cells were analyzed using a BD LSRFortessa. Populations were analyzed using FlowJo software.

### E1A expression and quantification of infectious progeny with AdNTE

To assess E1A expression 40,000 A549 cells were reverse transfected with siRNA and seeded into white 96-well plates and infected twenty-four hours later with AdNTE reporter virus at MOI 10. At end-point, media was removed and replaced with 20 uL room temperature serum free media and then 20 uL of Nano-Glo reagent (50:1 lysis buffer:Nano-Glo reagent). Cells were allowed to lyse for five minutes prior to quantification.

For quantifying infectious progeny cells were reverse transfected, seeded, and infected with AdNTE as previously mentioned into 96 well plates. At end-point plates were placed at -80°C, thawed, and then transferred to PCR strip tubes to perform an additional two freeze-thaw cycles. Infectious progeny was quantified in a second round of infection by seeding 40,000 A549 cells into white 96-well plates and infecting with 1/100 dilution of harvested virus. E1A expression determined by luminescence was quantified at 6 hpi as described above and the luminescent signal is proportional to infectious progeny production.

### Viral entry quantification with AdNV

Forty thousand A549 cells were reverse transfected with siRNA and seeded into white 96-well plates and infected twenty-four hours later with AdNV reporter virus at MOI 10. At twenty minutes post infection cells were washed 3x in PBS and luciferase activity was quantified as described above. A positive control was performed by pre-incubating AdNV virus with an anti-HAdV-C5 polyclonal antibody for 15 minutes prior to infection.

### Chromatin immunoprecipitation qPCR

Ten million A549 cells were infected with MOI 10 and cross-linked with 19 mL of 1% formaldehyde for ten minutes at room temperature. Cross-linking was quenched with the addition of 1 mL 2.5 M glycine for 5 minutes. Cells were washed 3x in ice cold PBS and harvested by scraping, cell pellets were snap frozen. Frozen cell pellets were thawed on ice and resuspended in 1 mL ChIP nuclear isolation buffer, incubated for 10 minutes on ice, and centrifuged at 1000g for 10 minutes at 4°C. Nuclei were lysed in 200 uL ChIP lysis buffer and sonicated for 8x 30 second ON/OFF cycles in a Bioruptor Pico. Sonicated samples were centrifuged to clear lysate and supernatant was diluted with 800uL ChIP dilution buffer. A 1% input was taken and sample was divided into corresponding number of IPs and incubated with antibody bound protein A dynabeads. Each IP used 15 uL dynabead and 3ug antibody. IPs were incubated overnight at 4°C with end-over-end rotation. The following day samples were washed once with ChIP wash buffer 1, twice with ChIP wash buffer 2, once with TE+0.1% Triton-X, and finally once with TE. Elution was carried out with ChIP Elution buffer for 30 minutes at room temperature with mixing on a thermomixer. Supernatant was transferred to new tubes containing 10 uL 5 M NaCl and 40ug Proteinase K, and incubated at 65°C for 4 hours. DNA was purified using the Zymo Research ChIP DNA clean and concentrator kit following the manufacturers protocol exactly. Immunoprecipitated DNA was quantified by qPCR using primers targeting the E1A promoter, E1A enhancer, tripartite leader, fiber open reading frame, and GAPDH (Primers and siRNA Table).

### Reverse transcription qPCR

A549 cells were reverse transfected with siEP400#1 and infected with wild type HAdV-C5 24 hours later. RNA was isolated from infected cells at 6 hpi using the Qiagen RNeasy kit following the manufacturer’s instructions. cDNA was prepared using the High-Capacity cDNA-to-RNA kit. Relative quantification of the major E1A transcripts (13S and 12S) and cellular (GAPDH) transcripts was performed using the Power SYBR Green PCR Master Mix. E1A mRNA was normalized to GAPDH and expressed as a relative fold change compared to siRNA control transfection.

### Quantification of viral genomes by qPCR

Viral genomes from infected cells were isolated using the PureLink Genomic DNA Mini Kit according to the manufacturer’s instructions. For relative quantification of incoming HAdV-C5 genomes in A549 cells DNA was harvested from infected cells at 4 hpi. Nuclear genomes were specifically collected using the REAP method^105^ prior to DNA extraction. For relative genome quantification in HDF-TERT cells following establishment of persistent infection, DNA was harvested 7 days post transfection with siEP400#1 or control siRNA. Viral DNA was quantified by qPCR with Power SYBR green using primers targeting the tripartite leader region and normalized to cellular GAPDH levels.

### Data analysis

Software used for data analysis can be found in the Reagents Table. Microscopy images were analyzed with ImageJ Fiji. All experiments in this study were performed with at least three biological replicates. Differential abundance analysis from whole cell proteome samples was performed with limma^106^ and identification of high confidence protein interactors was performed with the DEP package^107^ in R. For statistical analysis of protoemics data imputation was performed using a minimum probability distribution and only proteins quantified in more than 70% of samples were used for analysis. TcoF-DB^52^ was used to compare high confidence protein interactors with transcription regulators. For analysis of protein complexes the Cytoscape ClueGO plug-in^108^ was used to search Complex Portal^56^ and CORUM^55^. All other statistical tests were performed in GraphPad Prism.

## Acknowledgements

This manuscript is dedicated to Dr. Cason King who passed away in the early stages of this project. We are grateful to members of the Mymryk and Weitzman labs for discussion and feedback on the manuscript. We thank Dr. Patrick Hearing for providing HDF-TERT cells and guidance for the HAdV persistence model. We additionally thank Dr. Andrew Mehle for his valuable input. The authors thank Cassandra Wong and Brendon Seale of the Network Biology Collaborative Centre Proteomics Facility (RRID: SCR_025375) at the Lunenfeld-Tanenbaum Research Institute for proteomics services. The facility is supported by the Canada Foundation for Innovation and the Ontario Government. This research was partially funded by Canadian Institutes for Health Research (CIHR; PJT-2014110) grants and by grants to M.D.W. from the National Institutes of Health (R01 AI89552 and R01 AI118891). M.D.W. was supported in part by the Arthur Vincent Meigs Endowed Chair in Pediatrics. T.M.T. is partially supported by a CIHR post-doctoral fellowship.

## Author contributions

Conceptualization: T.M.T., J.S.M., and M.D.W. Methodology: T.M.T. Investigation: T.M.T., K.M.M., M.J.D., C.R.K., O.S.B., J.W.D. Writing – Original Draft: T.M.T. Writing - Review & Editing: T.M.T., J.S.M., M.D.W. Supervision: J.S.M. and M.D.W. Funding Acquisition: T.M.T., J.S.M., and M.D.W.

## Conflict of interest

None declared.

## Data availability

Proteomics data are available on the UCSD MassIVE repository (MSV000102285).

---

**Supplemental Figure 1.**
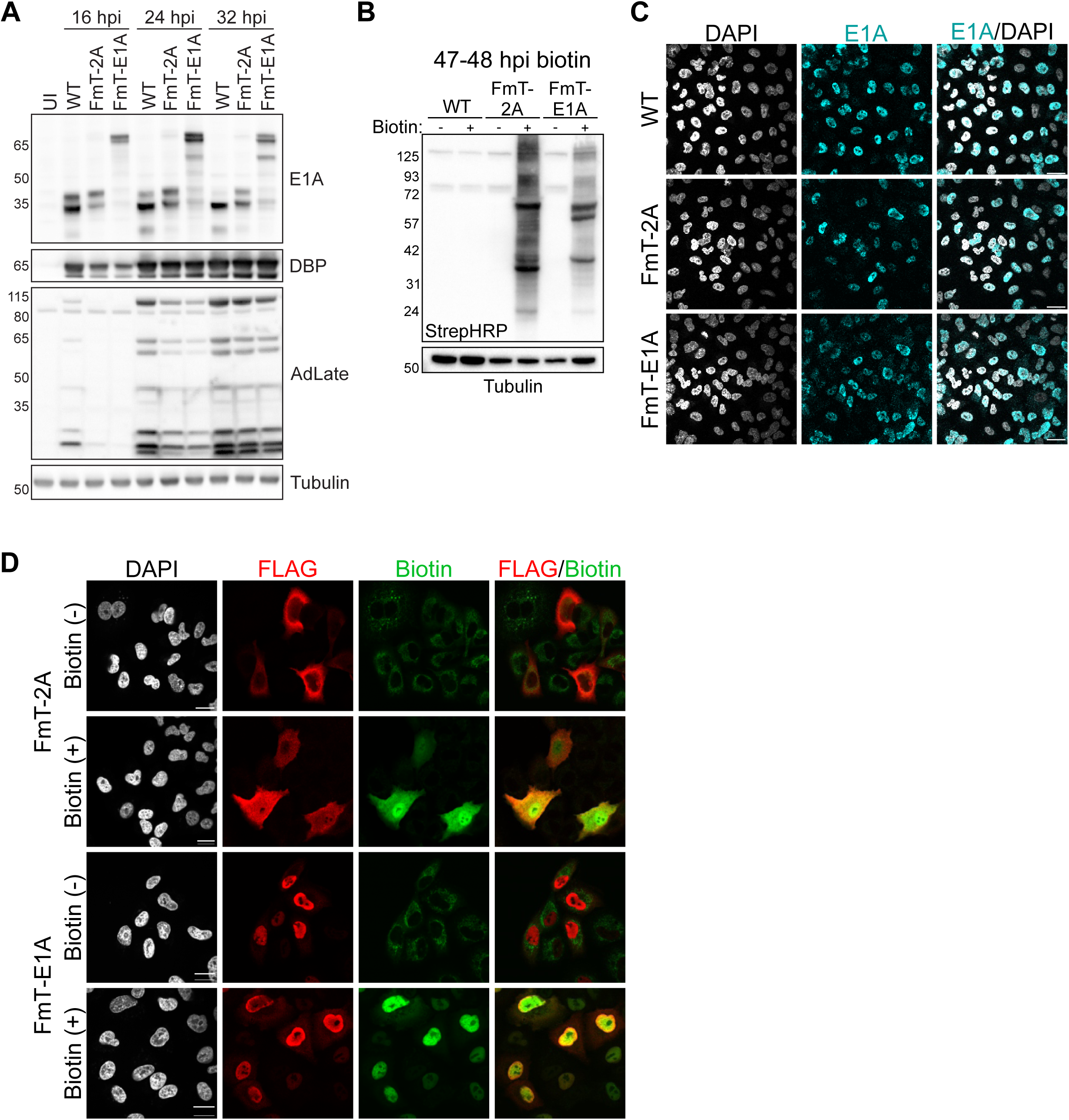
**A)** Western blot time course comparing wild-type HAdV-C5 and FmT viruses. Late protein expression was determined with a polyclonal antibody against HAdV-C5 capsid proteins (AdLate). **B)** Biotinylated protein in whole cell lysates from infected A549. Biotin was added for 1 hour at 47 hpi. **C and D)** Confocal immunofluorescence at 24 hpi in A549 cells infected with FmT-E1A or FmT-2A and immunostained for E1A (C), or FLAG and biotin (D). Scale = 20 µm.

**Supplemental Figure 2.**
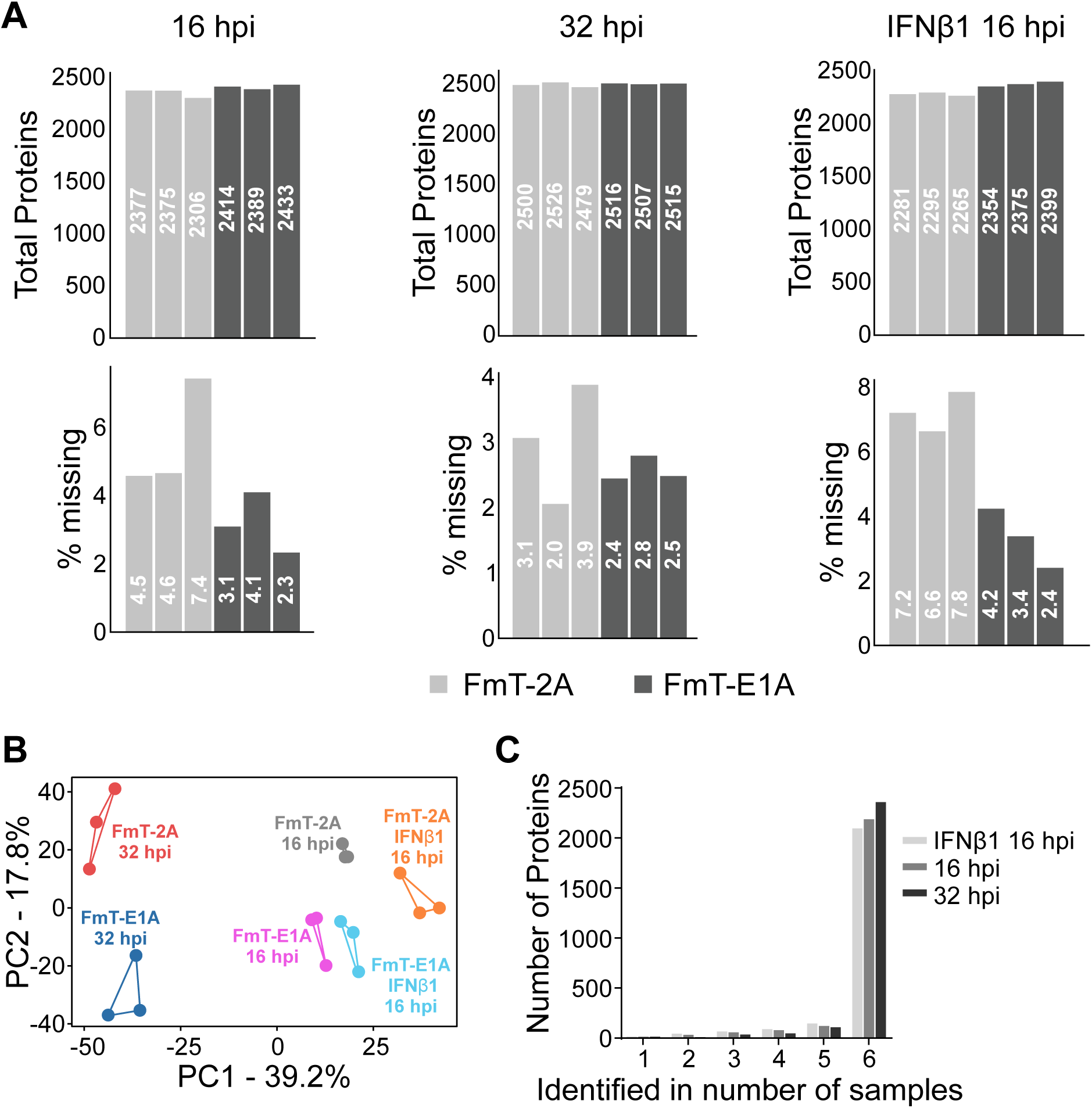
**A)** Quantified protein groups and corresponding percentage of missing values across samples and replicates. **B)** Principal component analysis. **C)** Reproducibility of proteins quantified in each condition.

**Supplemental Figure 3.**
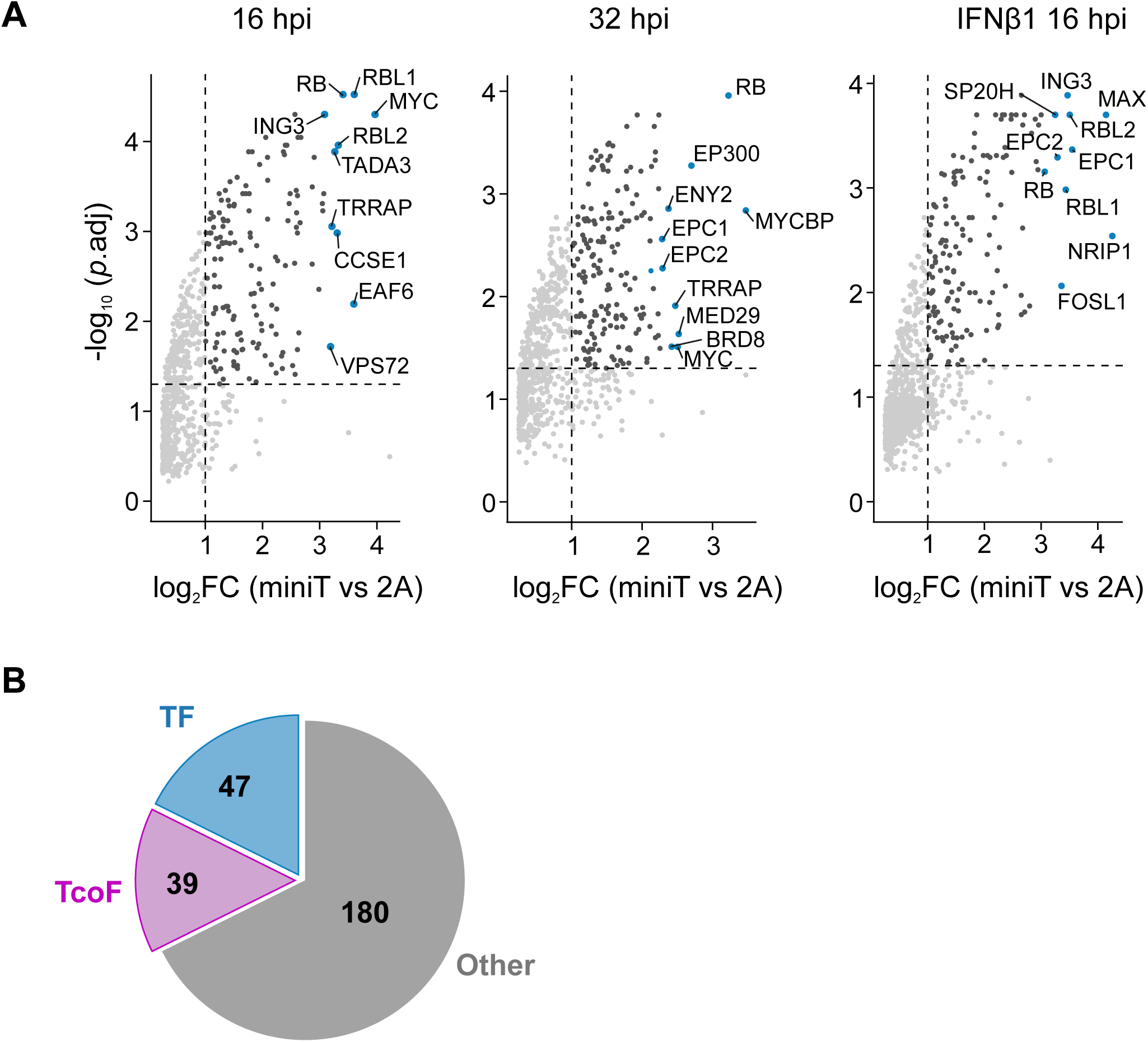
**A)** Volcano plots depicting high-confidence PPIs in each condition. Top 10 interactors based on log2 fold-change (FC) are labeled. **B)** Overlap of all high confidence interactors identified across conditions with the transcription (TF) and transcription cofactors (TcoF) in TcoF-DB.

**Supplemental Figure 4.**
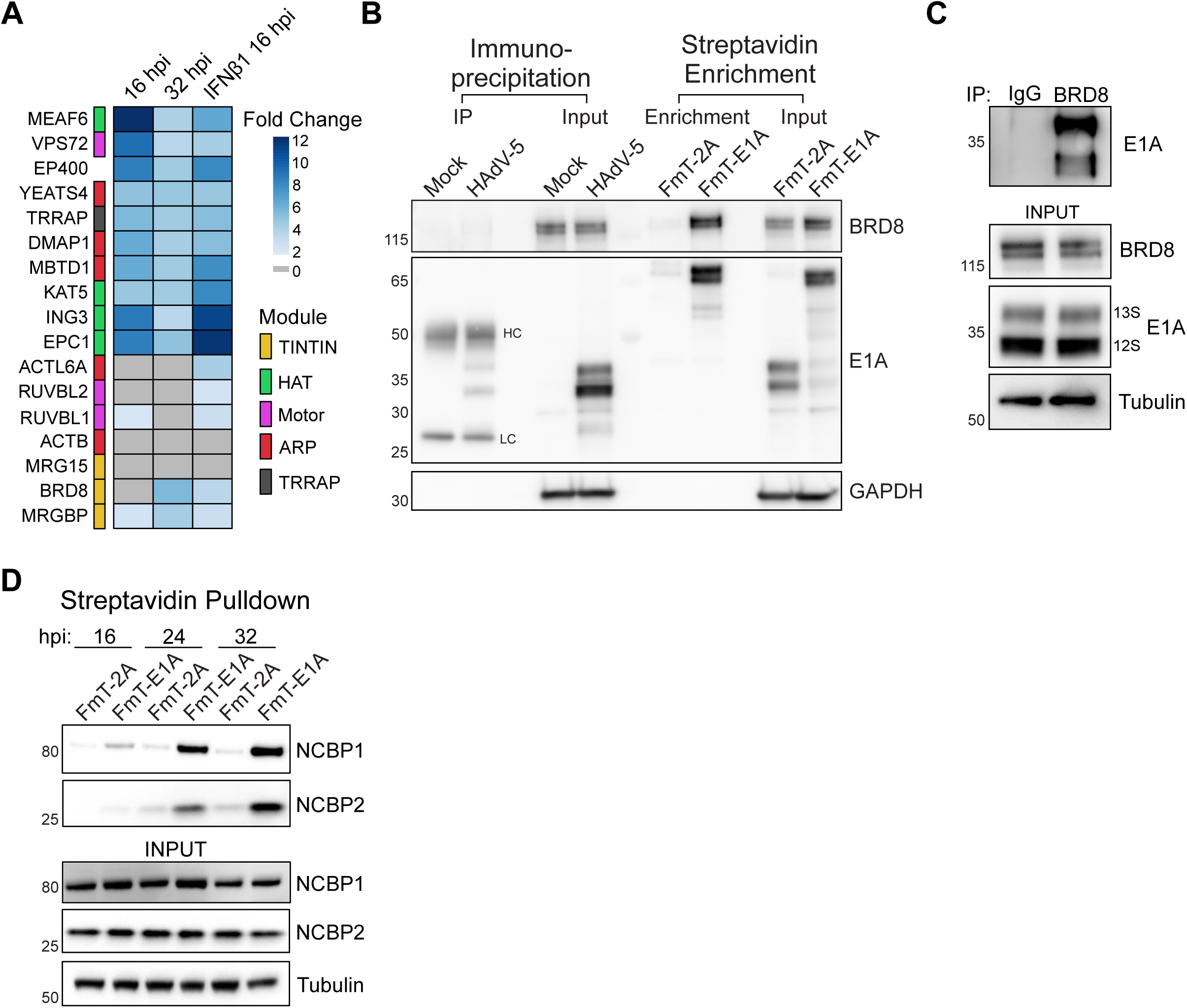
**A)** Heatmap depicting high-confidence interactors between E1A and p400 complex subunits. p400 subunits are colored based on which p400 module they belong to. Fold change is calculated as enrichment over the FmT-2A control. **B)** Comparison of E1A immunoprecipitation from HAdV-C5 infected A549 versus enrichment of biotinylated E1A interactors from cells infected with FmT-2A or FmT-E1A viruses. Both samples were harvested at 24 hpi and biotinylation was performed for 30 minutes prior to harvesting. HC = heavy chain, LC = light chain. **C)** BRD8 co-immunoprecipitation of E1A at 24 hpi in A549 cells infected with HAdV-C5. **D)** Time course of E1A-mediated proximity biotinylation of NCBP1/2 during infection with FmT-2A or FmT-E1A viruses. Cells were biotinylated for 30 minutes prior to harvesting and streptavidin enrichment.

**Supplemental Figure 5.**
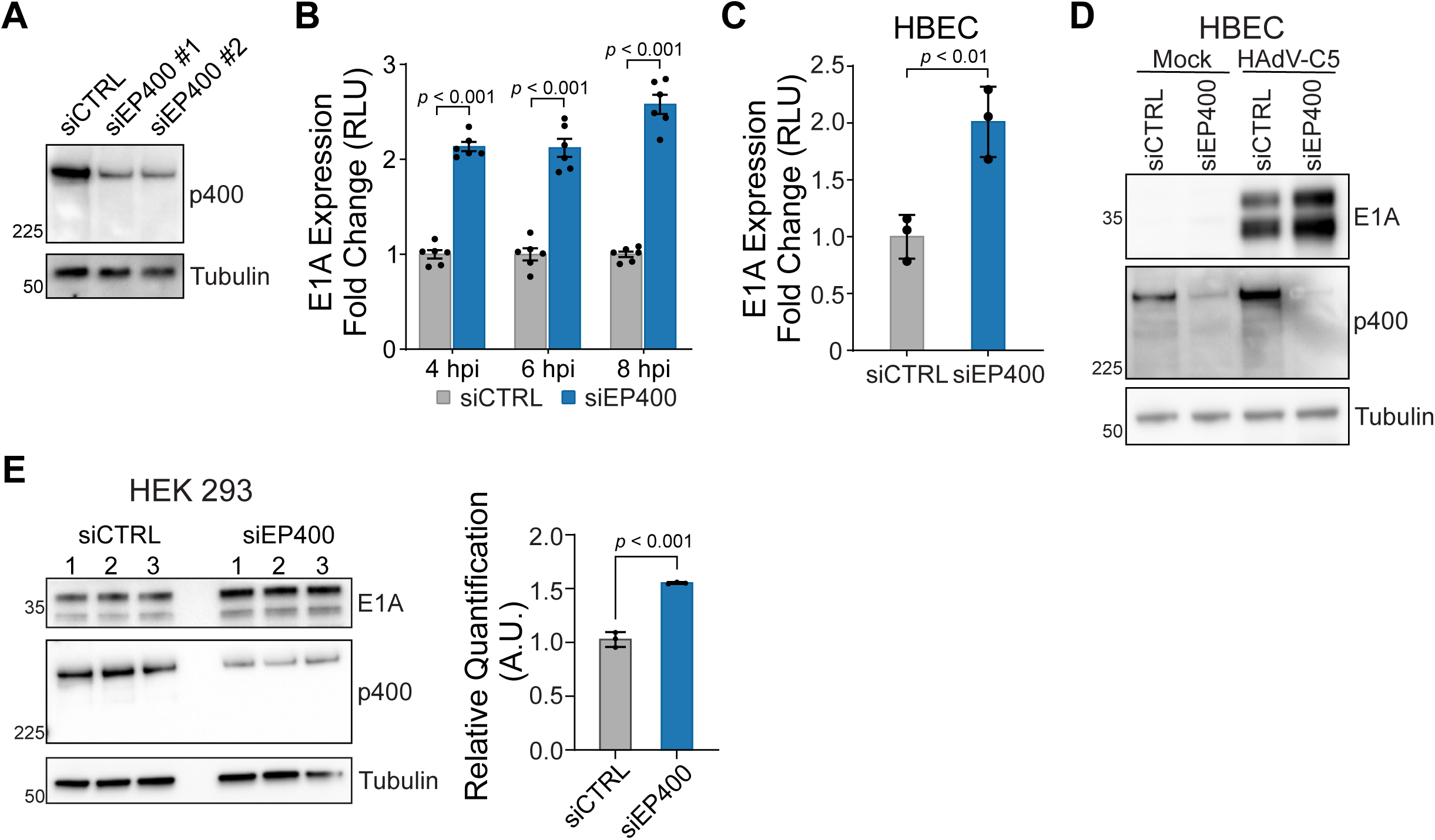
**A)** p400 siRNA test in A549 cells. p400 knockdown was determined via western blot at 24 hours post-transfection with siRNA. **B)** Relative quantification of E1A protein expression following p400 knockdown in A549 cells infected with AdNTE reporter virus. Time points correspond to Fig 4C. Each time point is normalized to its matched siCTRL. Multiple *t* test with Holm Sidak adjustment, n = 6. **C)** E1A protein expression in human bronchial epithelial cells (HBEC) following p400 knockdown. E1A expression was measured at 6 hpi with AdNTE reporter virus. *t* test, n = 3. **D)** E1A protein expression at 18 hpi in HBECs infected with HAdV-C5 following p400 knockdown. **E)** E1A protein expression in HEK 293 cells after 24 hours of p400 knockdown. Replicates were quantified in ImageJ and E1A was normalized to tubulin. *t* test, n = 3.

**Supplemental Figure 6.**
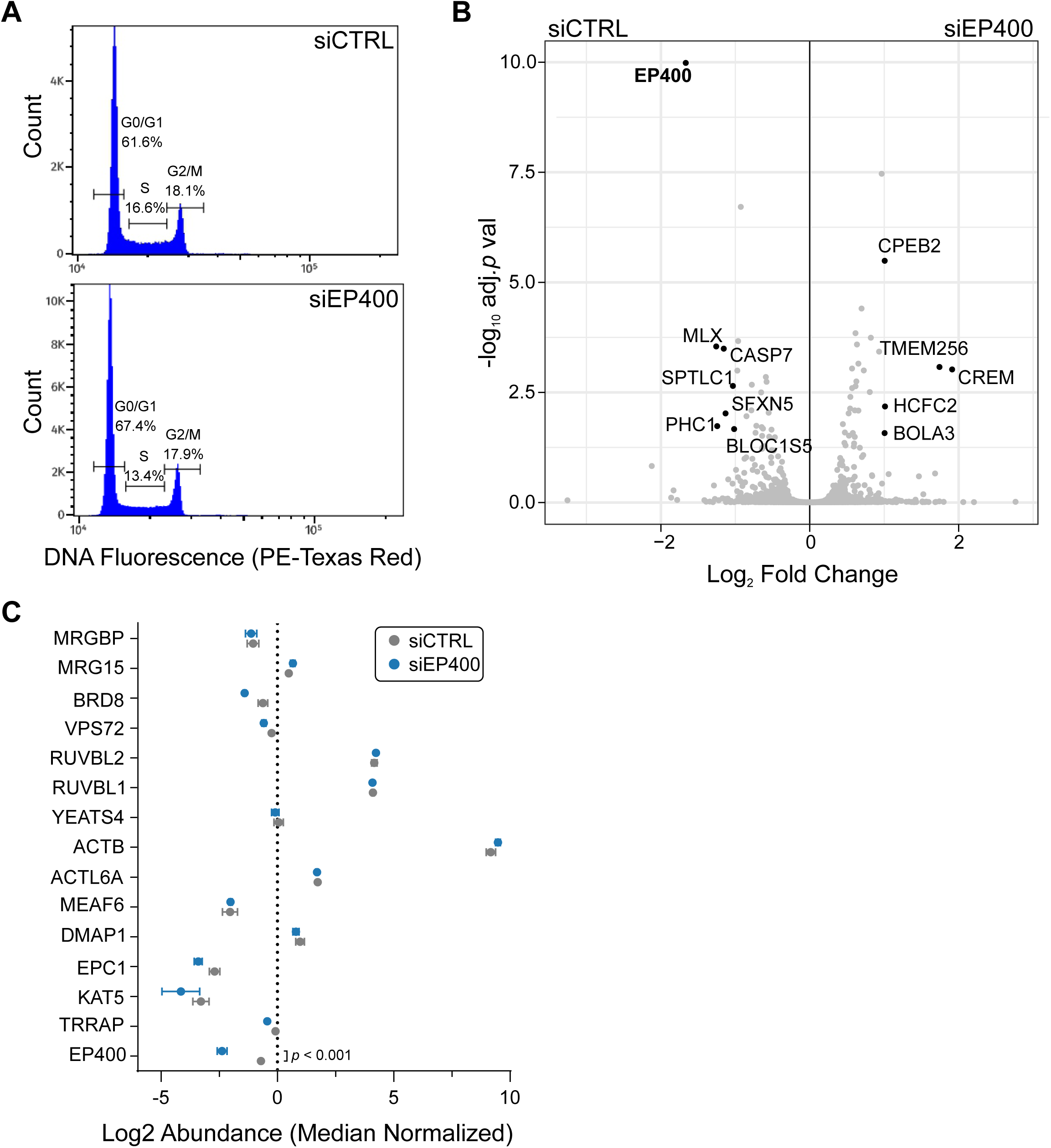
**A)** Flow cytometry analysis of cell cycle populations in uninfected cells following EP400 knockdown for 24 hours. **B)** Whole cell proteome analysis for differentially abundant proteins in uninfected A549 cells following p400 knockdown for 24 hours. Cutoffs are log2FC ≥ 1 and *p*.adj ≤ 0.05, n = 3. **C)** Protein abundance of p400 complex subunits quantified in whole cell proteome analysis. Only proteins with log2 fold-change ±1 and *p*.adj < 0.05 are displayed.

**Supplemental Figure 7.**
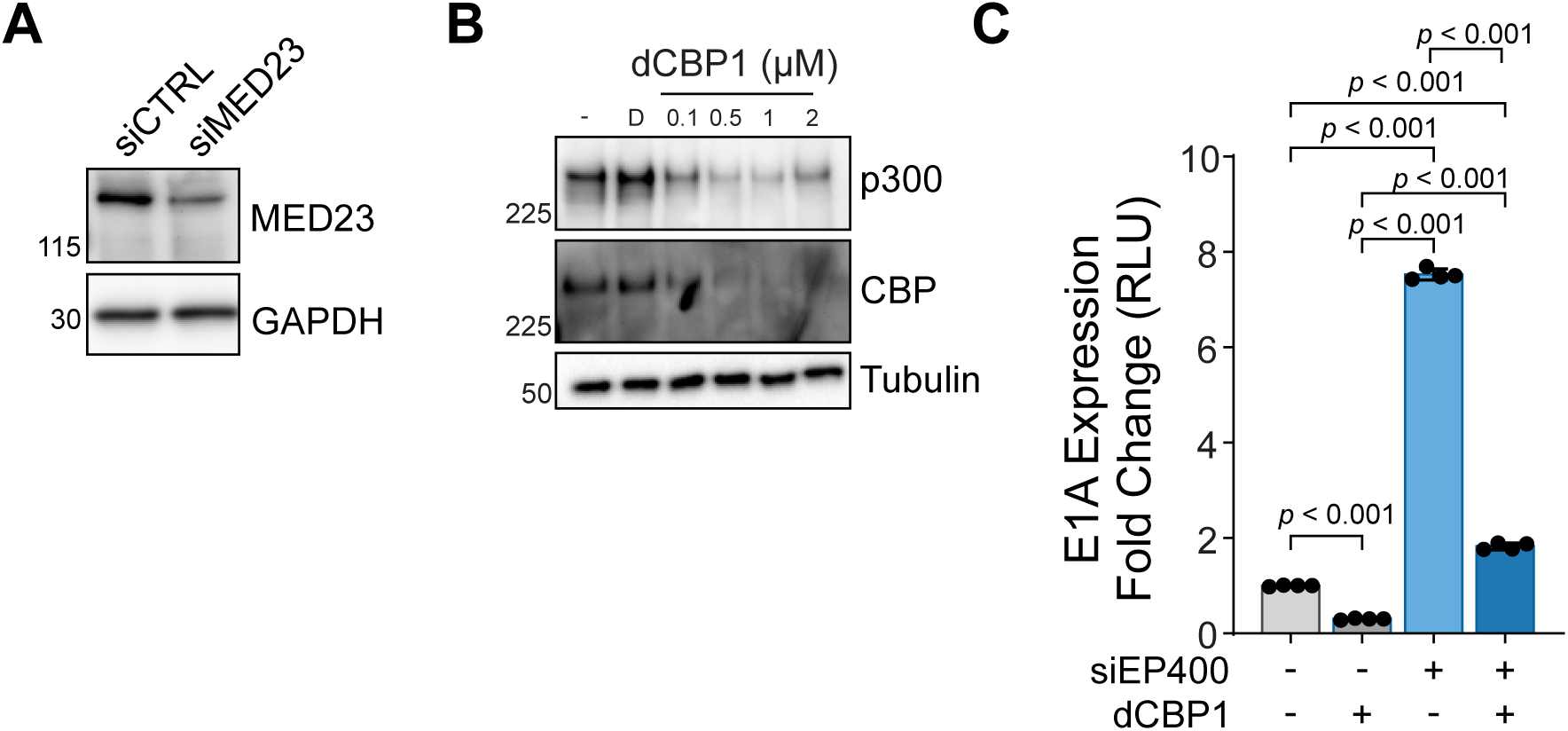
**A)** MED23 siRNA test in A549 cells. MED23 knockdown was determined via western blot at 24 hours post-transfection of siRNA. **B)** A549 cells treated with increasing p300/CBP degrader, dCBP1. Protein expression confirmed by western blot for p300 and CBP. “-“ = no drug, D = DMSO. **C)** E1A protein expression, measured by nanoluciferase activity, in cells infected with AdNTE reporter virus. Cells were transfected with siCTRL or siEP400 and treated with 1 µM dCBP1 prior to infection. E1A expression was assessed at 6 hpi. One-way ANOVA with Tukey’s correction, n = 4.

**Supplemental Figure 8.**
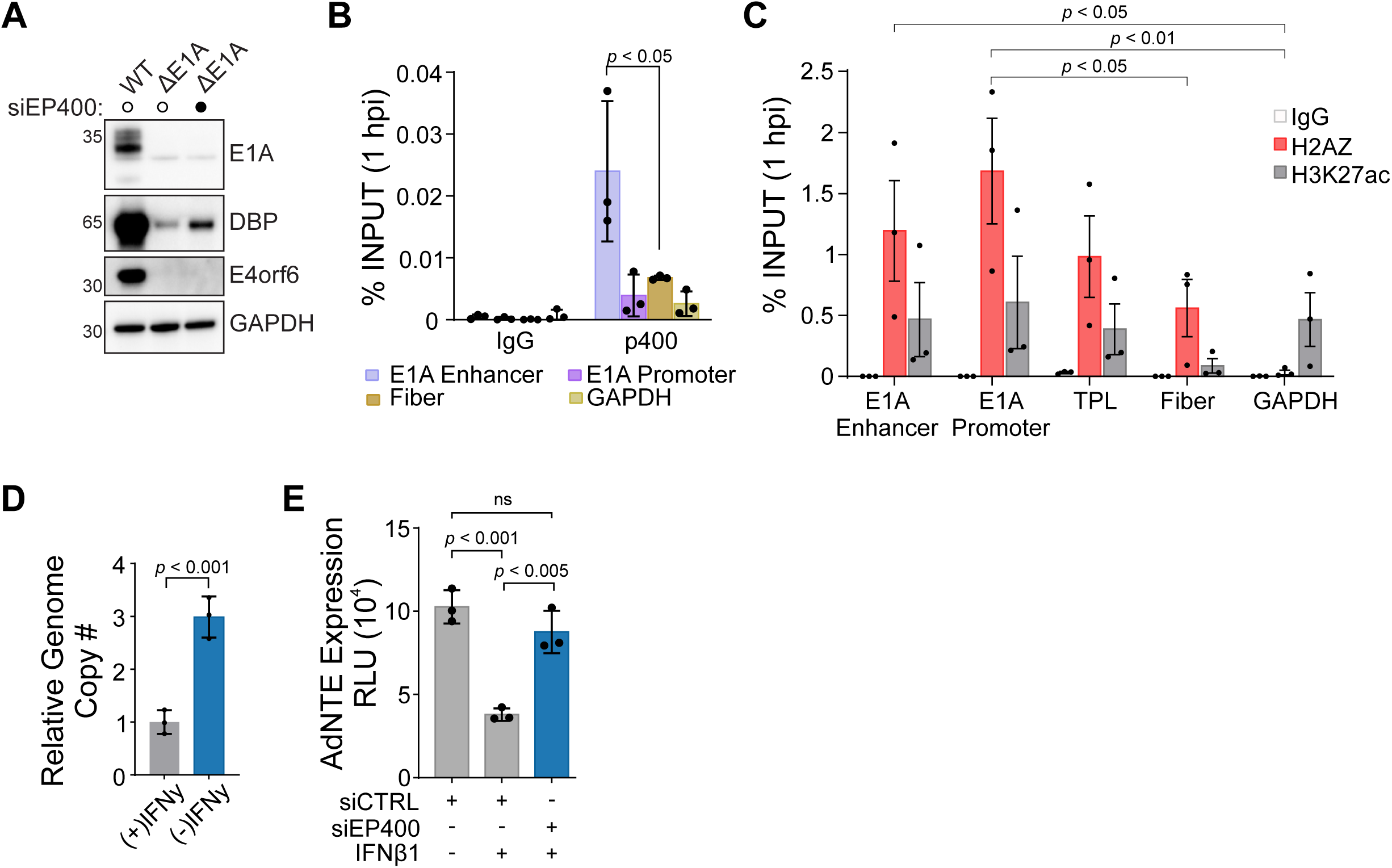
**A)** Viral early protein expression in A549 cells infected with WT HAdV-C5 or an E1A deleted virus (*dl309*) following p400 knockdown. WT infection with no siRNA control is used for reference viral protein expression. **B)** Association of p400 with HAdV-C5 genomes at 1 hpi. p400 ChIP-qPCR was performed at 1 hpi in A549 cells infected with HAdV-C5. Fiber was used to determine statistical enrichment of p400. One-way ANOVA with Fisher LSD, n = 3. **C)** Association of histone variant H2AZ with HAdV-C5 genomes at 1 hpi by ChIP-qPCR in A549 cells. GAPDH was used as a negative control for H2AZ and positive control H3K27ac. All statistical comparisons were made for H2AZ only. One-way ANOVA with Fisher LSD, n = 3. **D)** HAdVC-5 genome qPCR in TERT-HDF persistence model. At 30 days post infection IFNγ was removed and viral genomes were quantified 7 days post removal. *t* test, n = 3. **E)** E1A protein expression at 6 hpi following p400 knockdown and IFNβ1 treatment. E1A expression was measured by nanoluciferase activity in A549 cells infected with AdNTE reporter virus. One-way ANOVA with Tukey’s correction, n = 3.

## Notes

### Competing Interest Statement

The authors have declared no competing interest.

